# p75NTR and DR6 regulate distinct phases of axon degeneration demarcated by spheroid rupture

**DOI:** 10.1101/710111

**Authors:** Yu Yong, Kanchana Gamage, Irene Cheng, Kelly Barford, Anthony Spano, Bettina Winckler, Christopher Deppmann

## Abstract

The regressive events associated with trophic deprivation are critical for sculpting a functional nervous system. After nerve growth factor withdrawal, sympathetic axons maintain their structural integrity for roughly 18 hours (latent phase) followed by a rapid and near unison disassembly of axons over the next 3 hours (catastrophic phase). Here we examine the molecular basis by which axons transition from latent to catastrophic phases of degeneration following trophic withdrawal. Prior to catastrophic degeneration, we observed an increase in intra-axonal calcium. This calcium flux is accompanied by p75 neurotrophic factor receptor (NTR)-Rho-actin dependent expansion of calcium rich axonal spheroids that eventually rupture, releasing their contents to the extracellular space. Conditioned media derived from degenerating axons is capable of hastening transition into the catastrophic phase of degeneration. We also found that death receptor 6 (DR6) but not p75NTR is required for transition into the catastrophic phase in response to conditioned media but not for the intra-axonal calcium flux, spheroid formation, or rupture that occurs toward the end of latency. Our results support the existence of an inter-axonal degenerative signal that promotes catastrophic degeneration among trophically deprived axons.

## Introduction

Throughout nervous system development, axons, synapses and even entire neurons are initially overproduced and then refined through a period of organized culling (Purves and Lichtman, 1980; Hamburger and Oppenheim, 1990; Kantor and Kolodkin, 2003). Sympathetic neurons in the peripheral nervous system (PNS) rely on NGF-TrkA neurotrophic signaling to stabilize these components (Campenot, 1977; Purves et al., 1988; Hendry and Campbell, 1976; Yan et al., 2010; Gamage et al., 2017). The absence of sufficient neurotrophic signaling permits pro-destructive cues originating from tumor necrosis factor receptor superfamily (TNFRSF) members, such as p75 neurotrophin receptor (p75NTR), death receptor 6 (DR6) and tumor necrosis factor receptor 1a (TNFR1a), to promote regressive events like cell death, synapse restriction or axon degeneration in the PNS (Singh et al., 2008; Wheeler et al., 2014; Nikolaev et al., 2009; Olsen et al., 2014; Gamage et al., 2017). Axon degeneration is critical for proper nervous system wiring during development, but it’s also a hallmark of several neural pathologies like Alzheimer’s disease, amyotrophic lateral sclerosis (ALS), and injury (Saxena and Caroni, 2007). Both developmental and pathological degeneration occur in 2 phases: latent and catastrophic (Wang et al., 2012; Kristiansen and Ham, 2014).

The latent phase of degeneration commences immediately after neurotrophic factor deprivation and lasts roughly 18 hours *in vitro*. During this time, axons are morphologically indistinguishable from those receiving trophic support (Deckwerth and Johnson, 1993). Re-addition of trophic factor or mild depolarization during the latent phase is capable of rescuing degeneration (Edwards et al., 1991). After latency, the catastrophic/ execution stage of degeneration occurs in a rapid and near synchronous manner, taking roughly 3 hours to go from minimal to maximal degeneration amongst a population of axons *in vitro* (Gamage et al., 2017). During this period, axonal transport ceases, axons develop swellings, neurofilaments become fragmented, the cytoskeleton disintegrates, and debris is removed by recruited phagocytes (Saxena and Caroni, 2007; Luo and O’Leary, 2005). Activation of Bax-dependent apoptotic pathways and protease cascades involving calpain, caspase-9, -3, and -6 have been shown to promote catastrophic axon degeneration and commitment of cell death after NGF deprivation (Cusack et al., 2013; Simon et al., 2016, 2012; Ma et al., 2013; Yang et al., 2013). Importantly, this stage of degeneration represents the point of no return where re-introduction of neurotrophic factors do not rescue degeneration (Edwards et al., 1991). We and others have shown that in the absence of NGF-TrkA signaling, p75NTR and DR6 mediate axon degeneration (Singh et al., 2008; Olsen et al., 2014; Bamji et al., 1998; Nikolaev et al., 2009; Gamage et al., 2017). However, whether these receptors are involved in the latent or catastrophic phase of degeneration remains an open question. Given the apparent cooperativity of catastrophic degeneration amongst neurons, it stands to reason that degenerating axons might be communicating with one another, potentially through these death receptors.

How do axons transition from latent to catastrophic phases? It was shown that calcium signaling in degenerating axons promote activation of calpain, the calcium sensitive protease required for the disassembly of cytoskeletal elements (George et al., 1995; Avery et al., 2012; Vargas et al., 2015). A study in drosophila using calcium imaging revealed that compartmentalized calcium transients act as temporal and spatial cues to trigger dendrite pruning (Kanamori et al., 2013). Similar intra-axonal calcium waves have been observed prior to catastrophic fragmentation in injured zebrafish peripheral sensory axons (Vargas et al., 2015). Moreover, it was recently shown that axoplasmic calcium increases before the emergence of gross morphological changes in NGF deprived DRG cultures (Johnstone et al., 2019) suggesting that intra-axonal calcium signaling could play a role in all phases of degeneration.

We sought to identify the signaling events that occur during the transition between latent and catastrophic phases of degeneration induced by trophic withdrawal. To this end, we asked the following questions: **1.** What is the role of calcium in the latent and catastrophic phase in response to NGF deprivation? It may be that flux in intra-axonal calcium acts as a trigger for trophically deprived axons to exit the latent phase and enter the catastrophic phase. **2.** What are the signaling events that regulate the commitment to irreversible fragmentation? The engagement of calcium is well established in the execution phase of injury induced axon degeneration (Conforti et al., 2014), however whether other signaling pathways act permissively to allow calcium dependent irreversible fragmentation in response to trophic withdrawal remains an open question. **3.** What are the relative contributions of receptors p75NTR and DR6 to latent and catastrophic phases of degeneration? Because loss of p75NTR and DR6 showed different kinetics of axon degeneration after NGF deprivation (Gamage et al., 2017), we hypothesize that these death receptors may be required for different phases of degeneration.

Similar to injury paradigms, we demonstrate that after trophic deprivation, intra-axonal calcium increases prior to catastrophic degeneration. This is accompanied by the formation of calcium rich spheroids that grow and then rupture, releasing their contents (10 kDa and under) to the extracellular space while allowing an influx of extracellular molecules (*e.g.* calcium) into the intra-axonal space. We also found that pro-degenerative molecules released into the extracellular space are capable of hastening entry of trophically deprived axons into the catastrophic phase of degeneration. We next sought to define receptors that could initiate exit from latency or entry into the catastrophic phase of degeneration. We first examined whether p75NTR and/or DR6 were required for exit from latency by determining whether these receptors are required for spheroid formation and rupture. We found that p75NTR promotes spheroid formation, intra-axonal calcium rise, and membrane rupture in a Rho-dependent manner. In contrast, DR6 is not required for spheroid formation and rupture. We next asked whether these receptors promote entry into the catastrophic phase. We found that DR6 is required to hasten entry into the catastrophic phase of degeneration in response to conditioned media from degenerating axons, whereas p75NTR is dispensable. Taken together, these data place p75NTR and DR6 upstream and downstream of spheroidal rupture, respectively. This is consistent with separable roles for these receptors in early and late phases of degeneration induced by trophic withdrawal.

## Results

### Intra-axonal calcium increases in trophic factor deprived axons and accumulates in spheroids prior to catastrophic degeneration

To study the kinetics of degeneration induced by trophic withdrawal, we cultured sympathetic neurons from P0-P2 mice in microfluidic devices, which separate soma and axons (Fig.1a). After establishment in 45ng/mL of NGF for 5-7 days *in vitro* (DIV), wild-type sympathetic neurons were globally deprived of NGF with media containing anti-NGF function blocking antibody for indicated times. After NGF deprivation, the majority of axons remain intact for roughly 18 hours as measured by microtubule integrity (β3-tubulin staining). This period of sustained axonal integrity is referred to as the latent phase of degeneration (Coleman, 2005). At the conclusion of the latent phase, the majority of axons rapidly degenerate, going from 4.4±1.5% degeneration to 90.4±2.2% within 3 hours (Fig.1b-c). This is referred to as the catastrophic or execution phase of degeneration (Wang et al., 2012).

**Figure 1:**
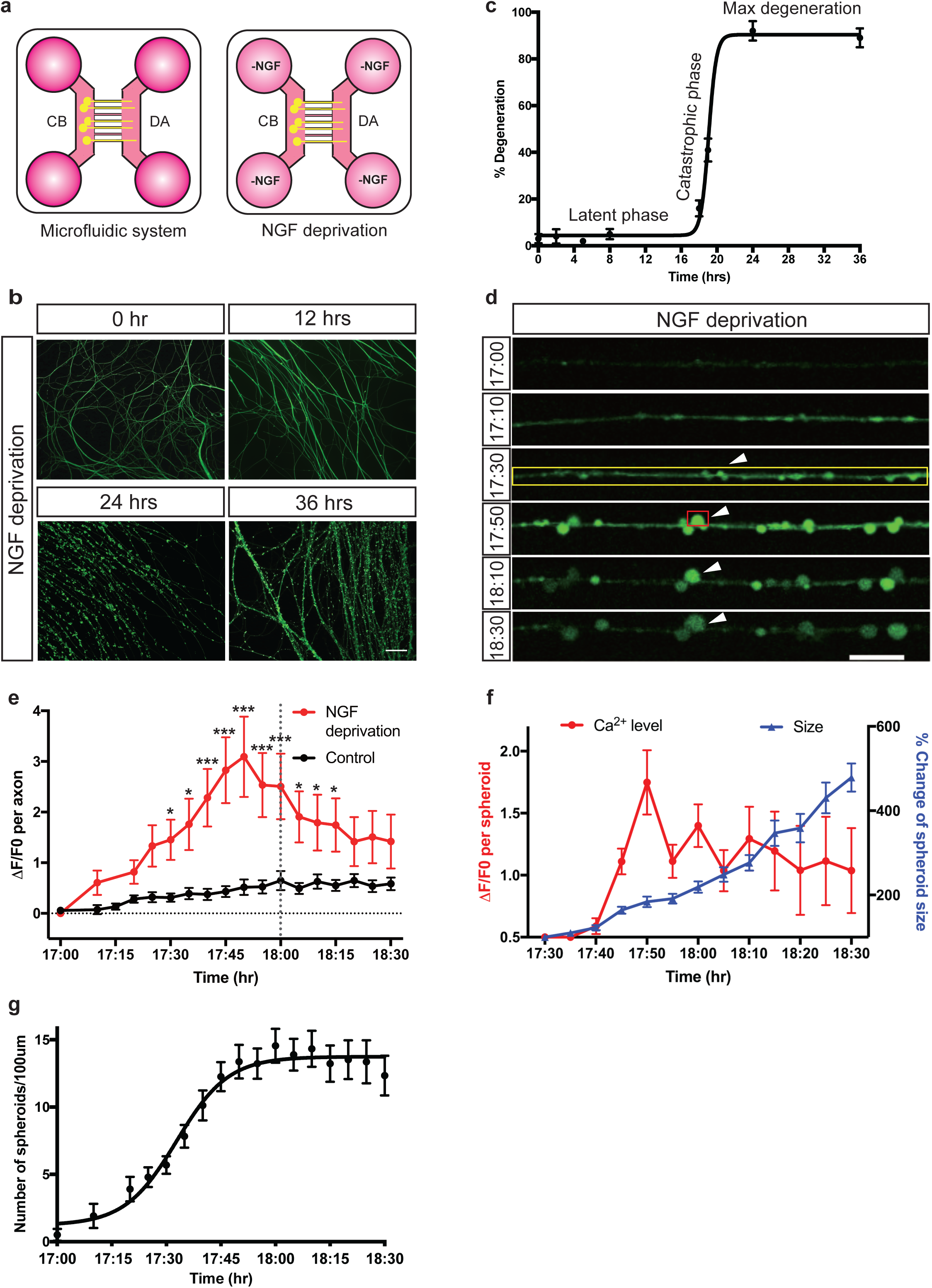
Axoplasmic calcium dynamics and formation of spheroids prior to catastrophic degeneration in response to NGF deprivation. a. Schematic representation of NGF deprivation paradigm in microfluidic devices. Cell bodies (CB) and distal axons (DA) are separated. For NGF deprivation, sympathetic neurons were maintained in NGF-deficient media containing 1µg/mL anti-NGF antibody. b. Representative images of β3-tubulin immuno-stained distal sympathetic axons before treatment (0hr), 12, 24 and 36 hours after NGF deprivation. Scale bar = 50µm. c. Degeneration time course after NGF deprivation. Latent and catastrophic phases of degeneration are noted. Nonlinear regression curves drawn according to Hill equation. n=3 for each time point. Data points represent as mean±SEM. d. Fluo4-AM calcium imaging of sympathetic axons at indicated times after NGF deprivation. Yellow box indicates individual axon as region of interest. Red box indicates axonal spheroid as region of interest. White arrowheads indicate the formation and growth of spheroid after 17 hours of NGF deprivation. Scale bar = 10µm. e. Calcium fluorescence change of control or NGF deprived axons over time. For the “NGF deprivation” condition, neurons were deprived of NGF for 17 hours, and then incubated with Fluo4-AM for calcium imaging. For the “Control” condition, no NGF deprivation was performed. Grey vertical dotted line indicates the onset of NGF deprivation induced catastrophic phase (18 hours after NGF deprivation). Black horizontal dotted line indicates the baseline without any calcium change. Total number of n=8 (NGF deprivation) and n=18 (control) of axons from 3 independent litters were quantified. Significant difference is determined by two-way ANOVA with multiple comparisons, *p<0.05, ***p<0.0001. Data shown as mean±SEM. f. Calcium fluorescence and size change of axonal spheroid after 17 hours of NGF deprivation. Total number of n=18 axonal spheroids from 3 independent litters were quantified. Data shown as mean±SEM. g. Quantification of axonal spheroid number per 100µm of axon at indicated times after NGF deprivation. Nonlinear regression curves were drawn according to Hill equation. Total number of n=11 axons from 3 independent litters were counted. Data shown as mean±SEM.

What are the molecular events that signal the transition from latent to catastrophic phase? Axonal calcium waves have been reported to be critical in promoting axon degeneration after axotomy (Vargas et al., 2015). Given the importance of the calcium dependent protease, calpain, in developmental axon degeneration, we speculated that calcium waves may occur at the transition between latent and catastrophic phases (Yang et al., 2013). We examined axonal calcium dynamics using Fluo4-AM imaging of axons that were withdrawn from NGF. Multiple small calcium peaks were detected during the first 12 hours, while the largest axoplasmic calcium flux that we observed occurred between 17-19 hours after trophic withdrawal (Supplementary Fig.1d), a time reflecting the transition window between latent and catastrophic phases (Fig.1d). Notably, axonal calcium increases up to 3 fold over baseline by 17hr50min after NGF deprivation (Fig.1e). After this initial increase, axonal calcium levels decrease approximately 1.5 fold. We hypothesize that axonal calcium increase may be a hallmark of the transition between latent and catastrophic degeneration phases.

In addition to axonal apoptosis induced by global NGF deprivation, pruning is critical for refinement of the nervous system during development. To examine whether axon pruning has similar calcium dynamics in relation to degeneration kinetics, we selectively deprived NGF in the axon compartment of sympathetic neurons grown in microfluidic devices (Supplementary Fig.1a). The majority of axons remain intact until 46 hours after local NGF deprivation, indicating the latent phase is significantly delayed compared to global NGF deprivation, which is consistent with previous reports (Chen et al., 2012) (Supplementary Fig.1b-c). Calcium levels in locally NGF deprived axons spiked prior to entry into the catastrophic phase of degeneration (Supplementary Fig.1d-e). This relationship between calcium spike and catastrophic degeneration is similar to what we observe in neurons globally deprived of NGF, albeit with an overall delay in degeneration kinetics.

Upon closer inspection of calcium dynamics within the entire axon, we observed that much of this calcium is concentrated in nascent spheroids on NGF deprived axons (Fig.1d, Supplementary Fig.1a). The formation of axonal spheroids/beads has been observed in not only trophic withdrawal models of developmental degeneration but also in pathological scenarios (Beirowski et al., 2010; Takahashi et al., 1997; Probst et al., 2000; Griffiths et al., 1998; Mejia Maza et al., 2018). However, whether these beads/spheroids have a function as degeneration progresses remains an open question. Interestingly, after observing the initial formation of calcium rich spheroids, these structures increase in size from 4.1±0.5μm^2^ to 19.6±2.7μm^2^ (roughly 500%) within one hour (Fig.1f). In addition to the increase in average spheroid size, the number of spheroids increased from 1 to 13 per 100μm between 17 and 18 hours after trophic withdrawal (Fig.1g). Spheroidal calcium levels had a roughly 1.7 fold increase by 17 hours and 50 minutes after NGF deprivation (Fig.1f). However, the spheroidal calcium levels varied depending on spheroid size. This value likely underestimates the increase in calcium since the spheroidal area measured increases with time. As such, we also normalized the areas used for our region of interest (ROI) to quantify the spheroidal calcium density and found the same trend (Supplementary Fig.1g).

To determine whether intra-axonal calcium elevation is required for the formation of spheroids, we depleted axoplasmic calcium by BAPTA-AM and examined the morphological changes of NGF deprived axons by live imaging. We find that axons retain their capacity to form spheroids in the absence of intra-axonal calcium elevation (Supplementary Fig.2a). These data indicate that while intra-axonal calcium flux may correlates with entry into the catastrophic phase of degeneration, it is not required for spheroid formation.

### Transcriptional induction of caspase activation is upstream of spheroid formation

Global NGF deprivation initiates apoptotic pathways to drive degeneration of axons and soma (Deckwerth and Johnson, 1993; Geden et al., 2019). Because caspases and calpain have been proposed to be involved in axon degeneration, we next examined the requirement of these proteases in the formation of spheroids after NGF deprivation. Application of the pan-caspase inhibitor Z-VAD-FMK but not calpain inhibitor III blocked spheroid formation (Fig.2a). Less than two spheroids per 100μm of axons were observed in NGF deprived cultures treated with Z-VAD-FMK, while the numbers of spheroids reached 10.2±0.4 and 8.8±0.2 per 100μm after 19 hours of NGF deprivation in the DMSO and calpain inhibitor III treatments, respectively (Fig.2b). Taken together, these data indicate that caspase activation is upstream of spheroid formation occurring at the transition between latent and catastrophic phases.

**Figure 2:**
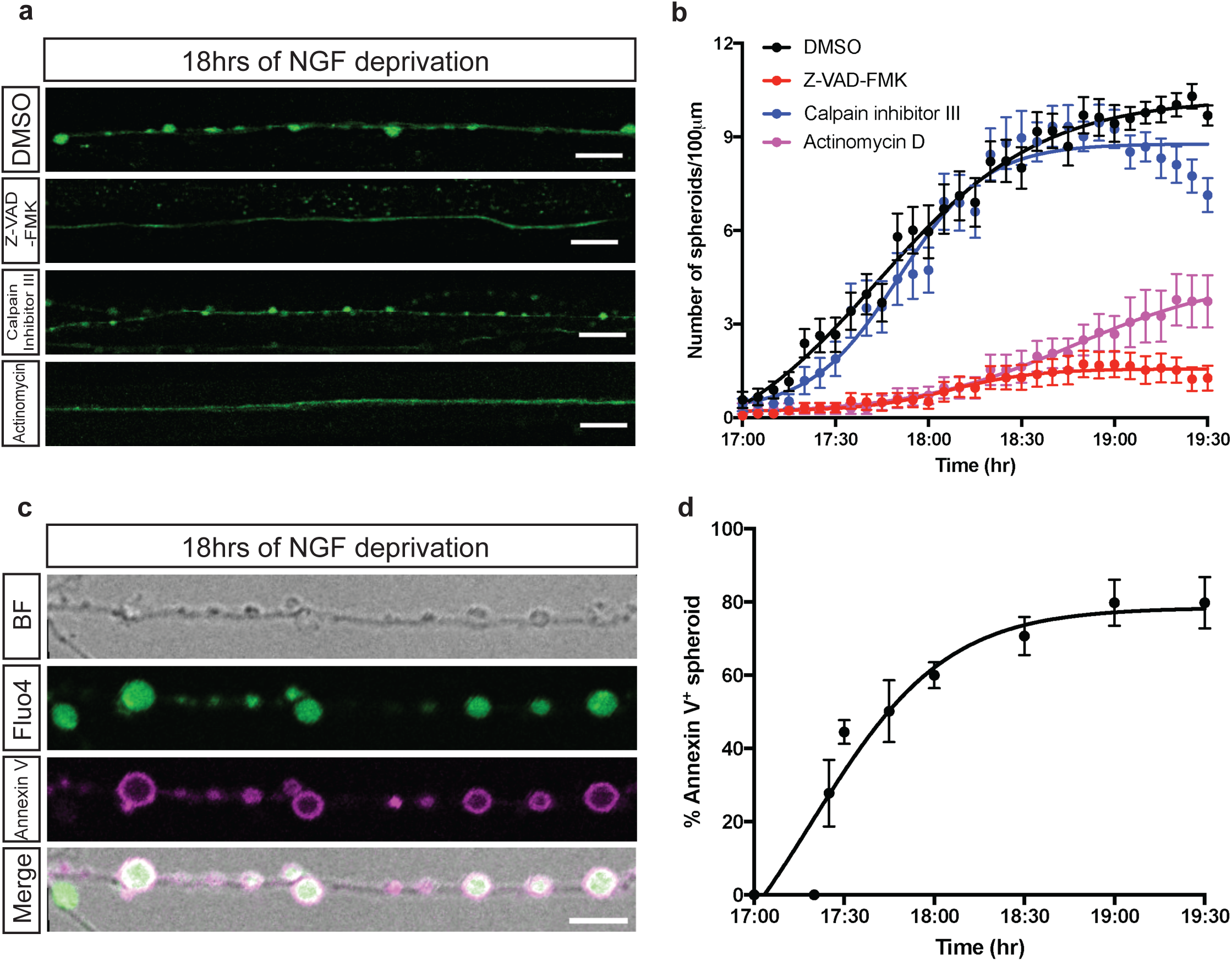
Transcription and caspase activation are required for formation of spheroids. a. Fluo4-AM calcium imaging of sympathetic axons after 18 hours of NGF deprivation in the presence of DMSO, 50µM V-ZAD-FMK, 20µM Calpain inhibitor III, and 1µg/mL Actinomycin, respectively. Scale bar = 10µm. b. Quantification of axonal spheroid number per 100µm of axons treated with DMSO (black), 50µM V-ZAD-FMK (red), 20µM Calpain inhibitor III (blue), or 1µg/mL Actinomycin (magenta) at indicated times after NGF deprivation. Nonlinear regression curves were drawn according to Hill equation. Total number of n=26 (DMSO), n=25 (Z-VAD-FMK), n=20 (Calpain inhibitor III), n=17 (Actinomycin) axons from 3 independent litters were counted. Data shown as mean±SEM. c. Representative images of axonal spheroids after 18 hours of NGF deprivation. Fluo4-AM (green) indicates intra-axonal calcium, and Annexin V (magenta) indicates exposure of phosphatidylserine on the extracellular surface of axonal spheroids. Scale bar = 5µm. d. Quantification of the percentage of fluorescent Annexin V positive spheroids after 17 to 19.5 hours of NGF deprivation. Total number of n=15 axons from 3 independent litters were counted. Data shown as mean±SEM.

It has been suggested that transcriptional upregulation of proapoptotic proteins (*e.g.* PUMA) are required for axon degeneration after trophic withdrawal (Simon et al., 2016; Maor-Nof et al., 2016). To determine whether inhibition of early prodegeneration transcriptional events could delay spheroid formation, we treated cell bodies with a transcription inhibitor, Actinomycin D, at the same time cells were deprived of NGF and examined spheroid formation between 17 and 19.5 hours after trophic withdrawal. Interestingly, Actinomycin D significantly diminished spheroid formation compared to the DMSO control (Fig.2a-b). Based on previous reports, we speculate that a transcriptional program induced by NGF withdrawal is permissive for the activation of caspase pathways, which in turn promotes spheroid formation (Fig.2a-b) (Simon et al., 2016; Maor-Nof et al., 2016).

Once cells become apoptotic, plasma membrane phosphatidylserine (PS) asymmetry is lost and membranes undergo blebbing. Importantly, PS enrichment on the outer leaflet of the plasma membrane is read as an ‘eat-me’ signal for phagocytes to aid apoptotic cell recognition and clearance (Poon et al., 2014; Zhang et al., 2018). After NGF deprivation, the formation of axonal spheroids were detected by phase-contrast imaging (Supplementary Fig.2b). We next assessed PS exposure on the extracellular surface of axonal spheroids by Annexin V staining, which fluoresces upon binding to PS (Fig.2c, Supplementary movie.1a,b). After 18 hours of NGF deprivation, about 60% of spheroids, but not the rest of the axon, are Annexin V-positive (Fig.2d, Supplementary Fig.2c), suggesting PS flipping to the outer leaflet on the spheroid membrane.

### Axonal spheroids develop membrane ruptures after NGF deprivation

Whereas Annexin V staining is consistent with PS flipping to the outer leaflet of the plasma membrane, it is also formally possible that the membrane integrity of growing spheroids is disrupted, allowing access of the 45 kDa Annexin V protein to the inner leaflet of the plasma membrane. To test this, we bathed control or NGF deprived axons in neutral fluorescent dextrans of different sizes (3, 10, 70 kDa). If there were any ruptures on the membrane after NGF deprivation, the dextran would immediately diffuse to the axoplasm (Fig.3a). The exclusion of dextran was maintained until 17 hours and 40 minutes after NGF deprivation, however by 17 hours and 50 minutes the axoplasm began to fill with 3 kDa fluorescent dextran (Fig.3b, Supplementary Movie.2a,c). The fraction of labeled spheroids decrease with increasing size of fluorescent dextran, and no significant spheroidal uptake was observed with 70 kDa dextran (Fig.3c), indicating that ruptures occur on a nano-scale and are permeable to molecules less than 10 kDa. Moreover, we observed non-punctate dextran positivity of spheroids simultaneously as Fluo4-AM dye was lost (Supplementary Fig.2d-e), consistent with membrane rupture. We do observe some spheroids that have punctate dextran labeling, consistent with macropinocytosis, however the events that we consider rupture are quite distinct and involve rapid and complete filling of the spheroid (Supplementary Movie 2b). If the labeling of Annexin V on spheroids was due to entry of the dye through membrane rupture, the number of spheroids filled with dextran of similar or smaller sizes should be no less than the number of Annexin V positive spheroids. However, only 6.5±3.1% of 10 kDa dextran positive spheroids were observed after 18 hours of NGF deprivation, significantly less than 45 kDa Annexin V labeled spheroids (Fig.2d,3c), suggesting that the flipping of PS occurs prior to the entry of dextran or membrane rupture.

**Figure 3:**
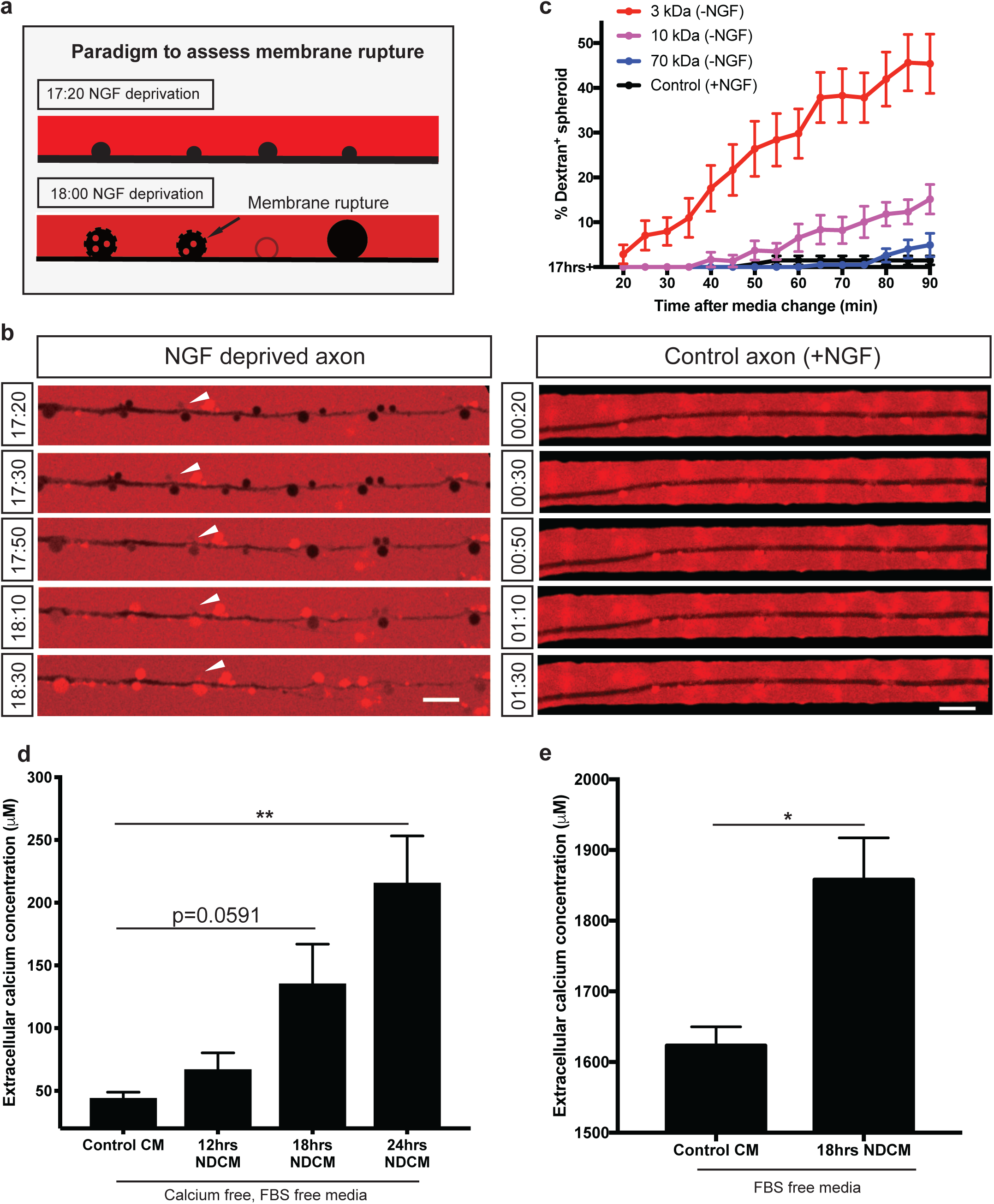
Axonal spheroids develop membrane rupture after NGF deprivation. a. Schematic representation of the experimental paradigm to assess membrane rupture model using fluorescent dextran. At 17 hours and 20 minutes of NGF deprivation, fluorescent dextran (red) is not taken up by the axon (black, negative space). However, by 18 hours of NGF deprivation, as the plasma membrane loses integrity and ruptures, fluorescent dextran (red) can diffuse into spheroids turning them red. Spheroids with intact membrane remain black. b. Representative images of dextran 3 kDa (red) entry to axonal spheroids (black) from 17 hours and 20 minutes to 18 hours and 30 minutes of NGF deprivation (left column), and dextran exclusion in untreated axons (right column). White arrowheads indicate that dextran 3 kDa enter to axonal spheroids after 18 hours of NGF deprivation. Scale bar = 10µm. c. Quantification of the percentages of fluorescent 3 kDa (red), 10 kDa (magenta), and 70 kDa (blue) dextran positive spheroids 20 to 90 minutes after 17 hours NGF deprivation. Black line (control) indicates the percentages of fluorescent 3 kDa dextran positive spheroids in the presence of NGF. Total number of n=15 (3 kDa), n=15 (10 kDa), n=12 (70 kDa), and n=9 (control) axons from 3 independent litters were counted. Data shown as mean±SEM. d. Measurement of extracellular calcium expelled from axons into calcium free, FBS free media. All axons were grow in regular DMEM media then switched to calcium free, FBS free media prior to NGF deprivation. In the “Control CM” group, media was collected from axons grown in the presence of NGF. In “NDCM” groups, medium were collected at 12, 18, and 24 hours after NGF deprivation. Values were analyzed from n=7 (Control CM), n=5 (12hrs NDCM), n=6 (18hrs NDCM), and n=7 (24hrs NDCM) independent replicates. Significance is determined by one-way ANOVA with multiple comparisons, **p<0.001. Data shown as mean±SEM. e. Measurement of extracellular calcium extruded from axons into FBS free, IS21 supplemented DMEM media with or without 18 hours of NGF deprivation. All axons were grow in FBS free, IS21 supplemented media prior to NGF deprivation. Values were analyzed from n=8 (Control CM) and n=3 (18hrs NDCM) independent replicates. Significance is determined by unpaired t-test, *p<0.05. Data shown as mean±SEM.

### Rupture of axonal spheroids disrupts the electrochemical gradient and allows expulsion of axoplasmic material

Just as membrane rupture allows free diffusion of molecules under 10 kDa to enter spheroids, we predicted that these ruptures would allow small molecules to freely diffuse out of spheroids. Indeed, once spheroids reach approximately 400% of their original size, we observe a diminution of calcium signal, which we speculate is a function of diffusion of Fluo4-AM and calcium out of the spheroid. We sought to use calcium release to the extracellular space as a surrogate to test the hypothesis that other small molecules may be released after spheroid rupture. We bathed axons in microfluidic devices in 100μl of calcium free media and measured extracellular calcium before and after spheroid formation using a Fluo4 spectrophotometric assay and CaCl_2_ standard curve (Supplementary Fig.3a). Sympathetic neurons were cultured in media containing FBS and 1.8mM calcium, then switched to calcium free, FBS free media. We found that 24hrs NDCM contained approximately 4-fold higher calcium (215.50±37.68μM) compared to Control CM (43.88±4.97μM) (Fig.3d). Additionally, we were also able to observe an increase in extracellular calcium after spheroid rupture when experiments were performed using regular culture media containing FBS and 1.8mM calcium. We found that at 18hrs NDCM displayed a roughly 50% increase in calcium (2.24±0.08mM) over Control CM (1.53±0.03mM) (Supplementary Fig.3b). To rule out the effect of FBS in regular culture media prior to calcium measurements, neurons were cultured in FBS free, IS21 supplemented DMEM media. Calcium level in 18hrs NDCM (1858.08±59.23mM) was significantly higher than calcium concentration in Control CM (1623.26±26.66mM) (Fig.3e), suggesting that this calcium release occurred regardless of the presence or absence of FBS in the media.

Beyond membrane rupture, it’s formally possible that calcium release could be in part due to enhanced permeability of plasma membrane Ca^2+^ ATPase (PMCA) and/or reversal Na^+^/Ca^2+^ exchanger (NCX) (Brini and Carafoli, 2011; Bano et al., 2007). To test this possibility, we treated NGF deprived axons with PMCA inhibitor caloxin 2A1 or NCX inhibitor bepridil to test their ability to block elevation of extracellular calcium concentration. Axons deprived of NGF for 18 hours and treated with caloxin 2A1 or bepridil retained their ability to release calcium to the extracellular space (Supplementary Fig.3c). Thus, spheroidal calcium is released to extracellular space independent on PMCA or NCX.

The observation that calcium in the media is elevated after membrane rupture is curious because the calcium concentration on the outside of the cell (1.8mM) is known to be much higher than in the axoplasm (100nM) (LaFerla, 2002; Clapham, 2007). How the axoplasm would have sufficient quantities of free calcium to appreciably elevate calcium levels in the media is unclear. To gain insight into this, we sought to determine the source of expelled intra-axonal calcium. We began by chelating all intracellular calcium with 10µM BAPTA-AM added 1 hour before NGF deprivation, which prevented calcium release to the extracellular space 18 hours after trophic deprivation (Supplementary Fig.3d). Intracellular calcium is buffered and stored by a contiguous network of ER and mitochondria, and calcium flux is thought to contribute to axon degeneration (Mattson, 2007; Villegas et al., 2014). Depletion of ER or mitochondria calcium, by treating neurons with 100nM thapsigargin or 20µM cyclosporine A for 12 hours prior to NGF deprivation also blocked calcium increase in the extracellular space (Supplementary Fig.3d).

We expect that in physiological scenarios, membrane rupture would also allow disruption of the calcium electrochemical gradient between the inside and outside of the cell. In this way, elevated calcium level in the extracellular space may represent a surrogate for the release of other small molecules as well as a sustained elevation of intracellular calcium, which may lead to activation of the calcium dependent protease, calpain. Importantly, we are unable to observe sustained elevation of axoplasmic calcium after rupture (Supplementary Fig.1d) because the calcium indicator, Fluo4-AM, used in the above experiments would also diffuse out of the axon.

### Expelled axoplasmic prodegenerative molecules hasten entry of axons into catastrophic phase

We speculated that contents released from ruptured spheroids may induce entry into the catastrophic phase of degeneration. To examine this, we collected media surrounding distal axons after 24 hours of NGF deprivation (24hrs NDCM) and incubated intact NGF deprived axons for 5 hours in 24hrs NDCM (Fig.4a). Importantly, in this paradigm the total time that recipient axons are without NGF is 17 hours, which is 1.5 hours prior to stereotypical morphological changes associated with degeneration. Neurons treated with control CM (from untreated axons at time 0 of NGF deprivation) displayed minimal degeneration, which is consistent with the kinetics of the trophic deprivation time course at 17 hours presented in Fig.1c. However, we found that intact axons treated with NDCM displayed 76.0±6.8% degeneration at this same time point (Fig.4b-c). Additionally, in the presence of NGF, neither NDCM nor elevated extracellular calcium promotes degeneration in 5 hours of incubation (Fig.4b-c), indicating that loss of NGF pro-survival signal is necessary to drive catastrophic axonal degeneration.

**Figure 4:**
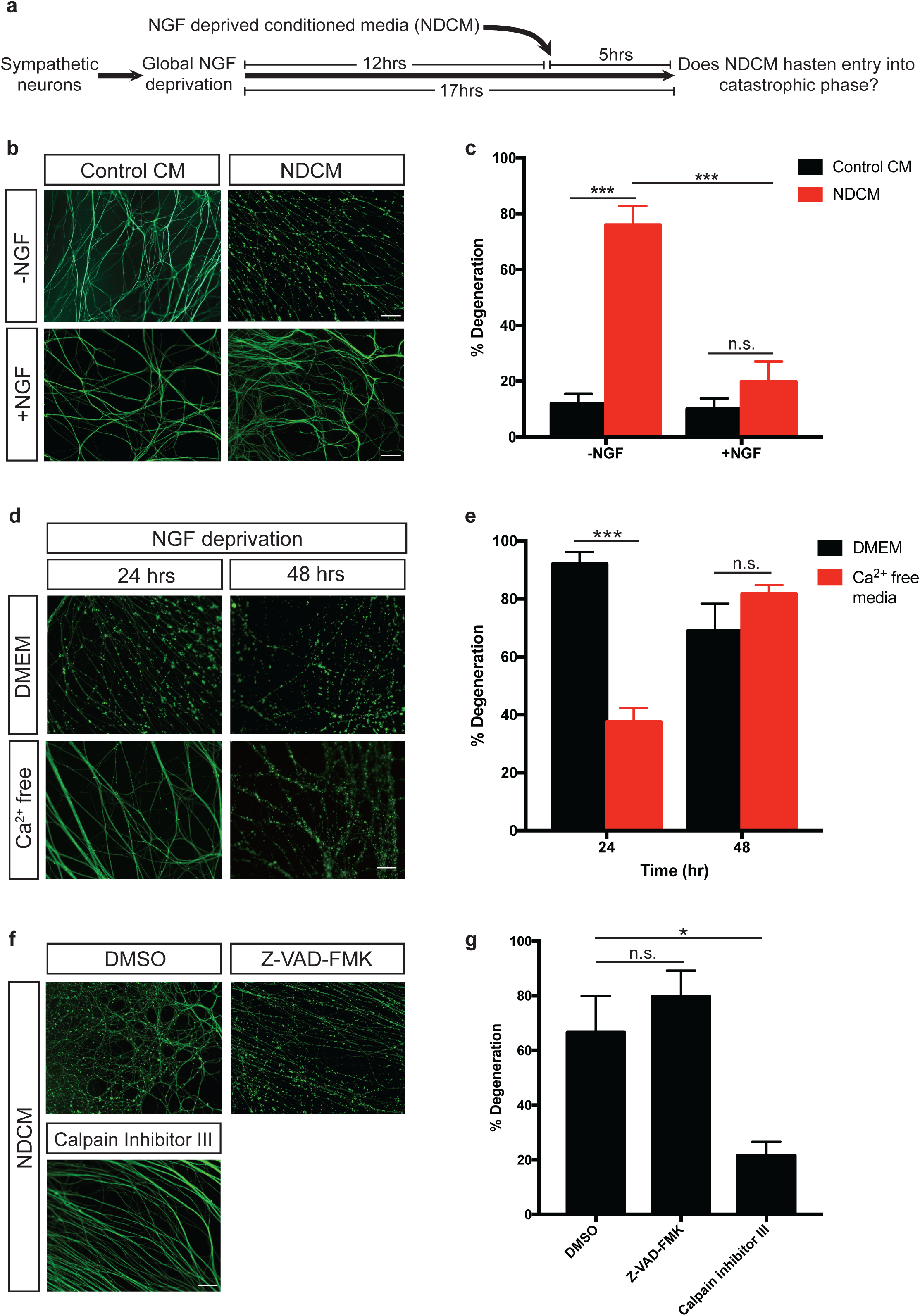
Calpain activation is required for NDCM-induced catastrophic axon degeneration. a. Conditioned media degeneration paradigm. WT sympathetic neurons were globally deprived of NGF for 12 hours followed by addition of conditioned media to the axons for 5 hours. NGF deprivation conditioned media (NDCM) was collected from degenerated axons after 24 hours of NGF deprivation. b. Representative images of β3-tubulin immuno-stained distal sympathetic axons after treatment with Control CM and NDCM for 5 hours in the presence and absence of NGF. Scale bar = 50 µm. c. Quantification of (b). Values are represented as mean±SEM. Significant difference is determined by two-way ANOVA with multiple comparisons, n.s.=not significant, ***p<0.0001. n=4 or more for each group. For each repeat, at least 100 axons were scored for degeneration. d. Representative images of β3-tubulin immunostained distal sympathetic axons after 24 and 48 hours of NGF deprivation in media with calcium (DMEM) and calcium free media. Axons were cultured in regular media (DMEM) and then switched to calcium free media at the time of NGF deprivation. Scale bar = 50 µm. e. Quantification of (d). Values are represented as mean±SEM. Significant difference is determined by two-way ANOVA with multiple comparisons, n.s.=not significant. ***p<0.0001, n=3 for each group. For each repeat, at least 100 axons were scored for degeneration. f. Representative images of β3-tubulin immunostained distal sympathetic axons after treatment with NDCM for 5 hours in the presence of DMSO, 50µM Z-VAD-FMK, and 20µM Calpain inhibitor III, respectively. All cultures were globally deprived of NGF for 12 hours prior to NDCM incubation. Scale bar = 50 µm. g. Quantification of (f). Values are represented as mean±SEM. Significant difference is determined by one-way ANOVA with multiple comparisons, n.s.=not significant, *p<0.05, n=7 or more for each group. For each repeat, at least 100 axons were scored for degeneration.

### Extracellular calcium and calpain activation are required for entry into the catastrophic phase of degeneration

Small pro-degenerative molecules may be released to extracellular space and re-enter axoplasma via membrane rupture to promote degeneration. If sustained elevation of intracellular calcium is required for axons to trigger catastrophic degeneration, we would expect that blocking the pro-degenerative positive feedback step by depleting extracellular calcium would delay catastrophic degeneration. To this end we performed trophic deprivation using calcium free DMEM. NGF deprived wild type axons maintained in DMEM with 1.85mM calcium, showed classic degenerative hallmarks like beading, blebbing and fragmentation by 24 hours (Fig.4d). Remarkably, when neurons were grown in calcium free media, their rate of degeneration slowed. After 24 hours of NGF deprivation only 38% of axons were degenerated in calcium free media, however maximal degeneration was observed 48 hours after NGF deprivation (Fig.4d-e). Thus, extracellular calcium is required to promote catastrophic degeneration in trophically deprived sympathetic axons. This is consistent with observations in trophically deprived sensory neurons and enucleated sympathetic neurons <u>(Johnstone et al. 2019; George et al. 1995)</u>.

To explore the downstream effector of calcium influx during developmental axon degeneration, we applied NDCM to recipient axons in the presence of calpain inhibitor III or Z-VAD-FMK. Interestingly, after 5 hours of incubation, only 21.6±5.0% of axons were degenerated in cultures treated with calpain inhibitor III, significantly less than the DMSO group (66.6±13.3%), while axons bathed in Z-VAD-FMK displayed 79.7±9.6% of degeneration (Fig.4f-g). These results indicate that calpain, instead of caspase, acts downstream of spheroid rupture and sustained calcium influx to promote catastrophic degeneration.

### p75NTR but not DR6 regulates the timing of spheroid formation and axoplasmic content expulsion to the extracellular space

Based on the above results, we have identified three events prior to catastrophic axon degeneration: **1.** An intra-axonal calcium increase; **2.** Calcium independent spheroid formation and membrane rupture; and **3.** Triggering of catastrophic degeneration by NDCM. What are the pathways upstream and downstream of these events? To address this we examined the role of two TNFR family members, DR6 and p75NTR, which we and others have implicated in axon degeneration following trophic withdrawal (Park et al., 2010; Twohig et al., 2011; Olsen et al., 2014; Gamage et al., 2017).

We first examined whether p75NTR or DR6 act upstream of spheroid formation and membrane rupture. To this end, we performed intra-axonal calcium imaging in NGF deprived *DR6*^*-/-*^ and *p75NTR*^*-/-*^ sympathetic axons. Both wild-type and *DR6*^*-/-*^ axons displayed up to a 5 fold increase of spheroidal calcium levels, however, *p75NTR*^*-/-*^ sympathetic axons did not display a significant increase in spheroidal calcium even after 20 hours and 30 minutes of NGF deprivation (Fig.5a-b). We next examined the role of p75NTR and DR6 in the accumulation of spheroids and the change in their size as a function of time after NGF deprivation. 18 hours after NGF deprivation, wild-type and *DR6*^*-/-*^ neurons display 12.0±1.0 and 6.1±0.7 spheroids per 100μm of axons, respectively (Fig.5a,c). However, *p75NTR*^*-/-*^ neurons displayed an approximately 2 hour delay in spheroid formation (Fig.5c), which is consistent with delayed onset of catastrophic degeneration (Gamage et al., 2017). We also observed that loss of *DR6* displays a roughly 800% increase in spheroid size after trophic deprivation similar to wild-type controls, whereas *p75NTR*^*-/-*^ neurons displayed a delayed accumulation of spheroids with a total size increase of less than 300% (Fig.5d). If spheroid formation is a prerequisite for membrane rupture, we would predict that NGF deprived neurons isolated from *p75NTR*^*-/-*^ mice would show diminished extracellular calcium expulsion whereas *DR6*^*-/-*^ would display normal calcium release, a surrogate for membrane rupture. Indeed, *p75NTR*^*-/-*^ axons showed no difference in extracellular calcium levels before or after 18 hours of NGF deprivation, while *DR6*^*-/-*^ neurons displayed a modest but significant increase (Supplementary Fig.3e). We next examined whether NGF deprived conditioned media from *p75NTR*^*-/-*^ or *DR6*^*-/-*^ neurons was capable of inducing degeneration of WT axons (Fig.5e). Consistent with calcium release phenotypes, we found that NDCM derived from *p75NTR*^*-/-*^ axons was incapable of inducing degeneration in WT axons (Fig.5d,h). In contrast, NDCM from *DR6*^*-/-*^ neurons contains prodegenerative activity (Fig.5g-h). Taken together, these data suggest that p75NTR but not DR6 is upstream of spheroid formation and membrane rupture.

**Figure 5:**
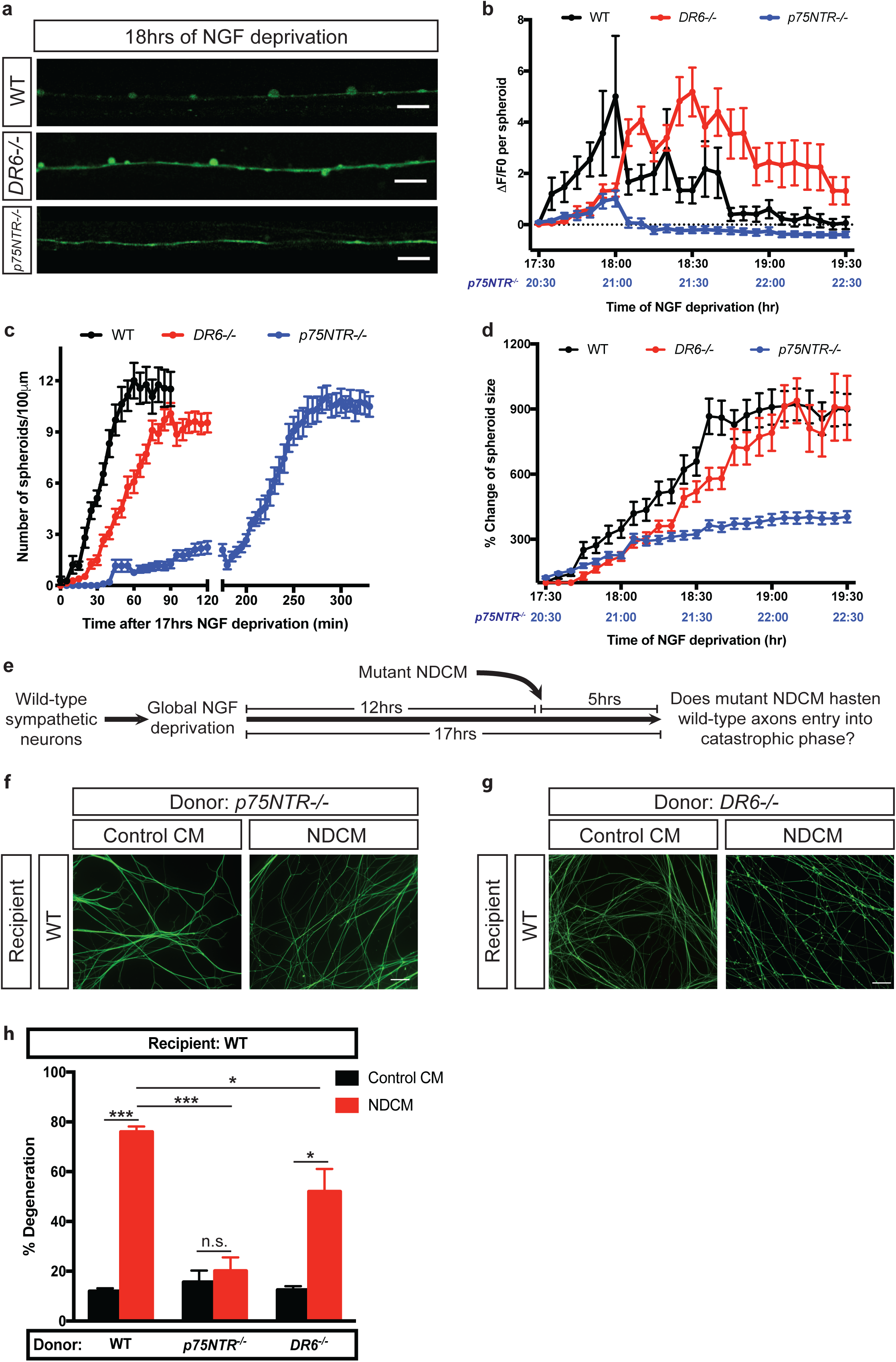
Depletion of p75NTR delays formation of spheroids and prodegenerative molecules exclusion after NGF deprivation. a. Fluo4-AM calcium imaging of wild-type, *DR6*^*-/-*^ and *p75NTR*^*-/-*^ sympathetic axons 18 hours of NGF deprivation. Scale bar = 10µm. b,d. Spheroidal calcium fluorescence (b) and size change (d) of wild-type, *DR6*^*-/-*^ and *p75NTR*^*-/-*^ sympathetic axons after indicated time of NGF deprivation. Black horizontal dotted line in (b) indicates the baseline without any calcium change. Lower x-axis labeled with blue correlates with time of NGF deprivation in *p75NTR*^*-/-*^ sympathetic axons. Individual axonal spheroids were quantified: n=32 (wild-type), n=30 (*DR6*^*-/-*^), and n=22 (*p75NTR*^*-/-*^) spheroids from cultured neurons harvested from 3 independent litters. Data shown as mean±SEM. c. Quantification of axonal spheroid number per 100µm of wild-type, *DR6*^*-/-*^ and *p75NTR*^*-/-*^ sympathetic axon at indicated times after 17 hours of NGF deprivation. Individual axonal spheroids were counted: n=20 (wild-type), n=29 ((*DR6*^*-/-*^), and n=22 (*p75NTR*^*-/-*^) axons from cultured neurons harvested from 3 independent litters. Data shown as mean±SEM. e. Catastrophic axon degeneration paradigm to test the pro-degenerative effect of mutant NDCM. Wild-type sympathetic neurons were globally deprived of NGF for 12 hours followed by addition of conditioned media derived from mutant axons for 5 hours. Mutant NDCM was collected from degenerating *p75NTR*^*-/-*^ or *DR6*^*-/-*^ axons 24 hours after NGF deprivation. f-g. Representative images of wild-type distal sympathetic axons immunostained for β3-tubulin after treatment with Control CM and NDCM collected from *p75NTR*^*-/-*^ (f) and *DR6*^*-/-*^ axons (g) for 5 hours. Scale bar = 50 µm. h. Quantification of (f-g).Values are represented as mean±SEM. Left two columns represent percentages of degeneration of wild-type sympathetic axons treated with wild-type NDCM and Control CM, respectively. Significance is determined by two-way ANOVA with multiple comparisons, n.s.=not significant, *p<0.05, **p<0.001, ***p<0.0001. n=3 or more for each group. For each repeat, at least 100 axons were scored for degeneration.

### p75NTR-Rho signaling is necessary and sufficient for spheroid formation and entry into catastrophic phase

How does p75NTR regulate spheroid formation prior to the catastrophic phase of degeneration? Previous work has shown that neurotrophins regulate growth cone dynamics by suppressing RhoA activation that occurs through p75NTR signaling (Gehler et al., 2004). p75NTR promotes RhoA activation to mediate sympathetic axonal degeneration in response to degenerative triggers like myelin (Park et al., 2010; Yamashita and Tohyama, 2003). Indeed, active Rho in axonal spheroids was detected by incubation with GST-rhotekin Rho binding domain (RBD) fusion protein after 18 hours of NGF deprivation but not in cultures treated with Rho inhibitor C3 transferase (CT04) (Fig.6a). Moreover, inhibiting Rho family members using CT04 (1µg/mL, 2 hours prior to 17 hours NGF deprivation) prevented spheroid formation in wild-type axons (Fig.6b-c). Consistent with this, axons treated with CT04 delayed degeneration for up to 24 hours after NGF deprivation (Fig.6d-e). The major downstream target of Rho activation is actin remodeling, and we similar to previous observations we observe actin and β3-tubulin enrichment in axonal spheroids (Supplementary Fig.4a) (Beirowski et al., 2010). However, studies have shown that Taxol, which stabilizes microtubules did not prevent spheroid formation, suggesting a secondary role of microtubule disruption (Barsukova et al., 2012). We next tested whether actin remodeling is required for spheroid formation. Indeed the actin polymerization inhibitor cytochalasin D (10µg/mL) inhibited spheroid formation and axon degeneration (Fig.6b-e, Supplementary Fig.4b). Taken together, these data suggest that activation of Rho and actin remodeling is necessary for spheroid formation and entry into the catastrophic phase of degeneration. Another downstream effector of Rho is the Rho-associated protein kinase 1 (ROCK1). During apoptosis, ROCK1 has been shown to be cleaved by caspase-3, resulting in constitutive kinase activity to promote phosphorylation of myosin light chain (MLC) and membrane blebbing (Sebbagh et al., 2001; Coleman et al., 2001). This may explain our finding of suppressed spheroid formation after NGF deprivation in the presence of caspase inhibitor (Fig.2a-b).

**Figure 6:**
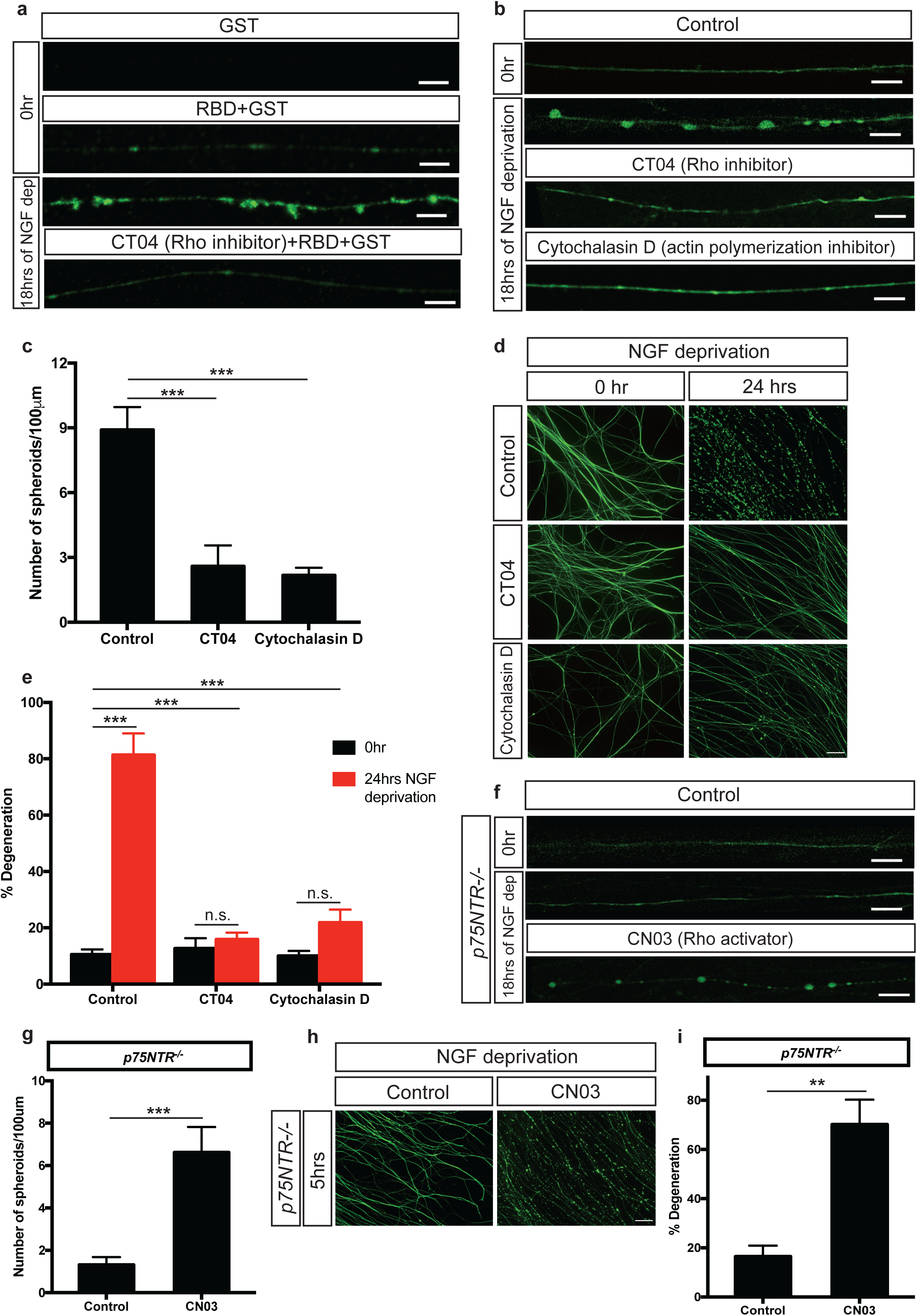
p75NTR-Rho signaling is required for axonal spheroid formation. a. Representative images of wild-type sympathetic axons immunostained for GST tag with or without NGF deprivation. All images except the first one show axons incubated with Rhotekin-RBD GST-fusion protein that binds active Rho proteins after fixation. Axon in the bottom image was NGF deprived and treated with 1µg/mL Rho inhibitor CT04 for 3 hours. Scale bar = 5 µm. b. Fluo4-AM calcium imaging of wild-type grown in NGF or NGF deprived sympathetic axons with or without drug treatment. For the “CT04” group, wild-type axons were incubated in SCG media containing 1µg/mL Rho inhibitor CT04, for 2 hours prior to 17 hours of NGF deprivation. For the “Cytochalasin D” group, wild-type axons were incubated in SCG media containing 10µg/mL actin polymerization inhibitor for 2 hours prior to 17 hours of NGF deprivation. Scale bar = 10µm. c. Quantification of axonal spheroid number per 100µm of wild-type sympathetic axons after 18 hours of NGF deprivation with or without CT04 or Cytochalasin D treatment. Total number of n=9 (Control), n=8 (CT04), n=22 (Cytochalasin D) axons from cultured neurons harvested from 3 independent litters were quantified. Significant difference is determined by ordinary one-way ANOVA with multiple comparisons, ***p<0.0001. Data shown as mean±SEM. d. Representative images of wild-type distal sympathetic axons immuno-stained for β3-tubulin with and without and 24 hours after NGF deprivation in the absence and presence of CT04 or Cytochalasin D. Scale bar = 50µm. e. Quantification of (d). Values are represented as mean±SEM. Significant difference is determined by two-way ANOVA with multiple comparisons, ***p<0.0001, n.s.=not significant, n=3 or more for each group. For each repeat, at least 100 axons were scored for degeneration. f. Fluo4-AM calcium imaging of *p75NTR*^*-/-*^ sympathetic axons grown in the presence or absence of NGF with or without CN03 treatment. For “CN03” group, *p75NTR*^*-/-*^ axons were incubated in SCG media containing 1µg/mL Rho activator CN03 for 2 hours before 17 hours of NGF deprivation. Scale bar = 10µm. g. Quantification of number of spheroids per 100µm of *p75NTR*^*-/-*^ sympathetic axons after 18 hours of NGF deprivation in the absence and presence of CN03. Individual axons were counted: n=16 (Control) and n=12 (CN03) axons from 3 independent replicates. Significance is determined by unpaired t test, ***p<0.0001. Data shown as mean±SEM. h. Representative images of *p75NTR*^*-/-*^ distal sympathetic axons immuno-stained for β3-tubulin after treatment with or without CN03 for 5 hours. All cell cultures were pre-treated with 12 hours of NGF deprivation. Scale bar = 50µm. i. Quantification of (h). Values are represented as mean±SEM. Significance is determined by unpaired t test, **p<0.001, n=7 (Control) and n=3 (CN03). For each repeat, at least 100 axons were scored for degeneration.

If Rho is downstream of p75NTR, we would predict that ectopic activation of Rho would promote spheroid formation and axon degeneration in the absence of *p75NTR*. To test this, we incubated NGF deprived *p75NTR*^*-/-*^ axons with Rho activator, CN03 (1µg/mL, 2 hours prior to 17 hours NGF deprivation). CN03 rescued the diminished spheroid formation and extracellular calcium release through membrane rupture in *p75NTR*^*-/-*^ axons 18 hours after NGF deprivation (Fig.6f-g, Supplementary Fig.4c-d). We also found that CN03 could rescue the timing of degeneration in the absence of *p75NTR* (NGF deprived for 12 hours with CN03 for the final 5 hours) (Fig.6h-i). Taken together these data suggest that p75NTR-Rho signaling is necessary and sufficient to promote spheroid formation and entry into the catastrophic phase of degeneration.

### p75NTR dependent spheroid formation is ligand dependent

We next sought to determine whether p75NTR requires ligand to induce spheroid formation. Application of LM11A-31, a small nonpeptide molecule competitor of p75NTR ligand binding (Massa et al., 2006; Simmons et al., 2014), suppressed spheroid formation on wild-type sympathetic axons after NGF deprivation (Supplementary Fig.5a-b). However, blocking brain-derived neurotrophic factor (BDNF), which has been shown to activate p75NTR apoptotic signaling (Kohn et al., 1999), failed to inhibit development of spheroids on NGF deprived axons (Supplementary Fig.5a-b). These results suggest that p75NTR mediated spheroid formation can be activated by binding pro-degenerative ligands other than BDNF.

### DR6, but not p75NTR, gates entry into the catastrophic phase of degeneration in response to NDCM

We next asked whether DR6 or p75NTR are downstream of the prodegnerative activity of NDCM using the paradigm described in Fig.4a. We applied NDCM collected from wild-type axons to recipient neurons derived from *DR6*^*-/-*^ and *p75NTR*^*-/-*^ mice. After a 5 hour treatment of NDCM, *p75NTR*^*-/-*^ axons showed 71.6±5.8% degeneration, while *DR6*^*-/-*^ axons displayed minimal degeneration (Fig.7a-b). These data suggest that DR6 but not p75NTR is downstream of the prodegenerative effects of NDCM.

**Figure 7:**
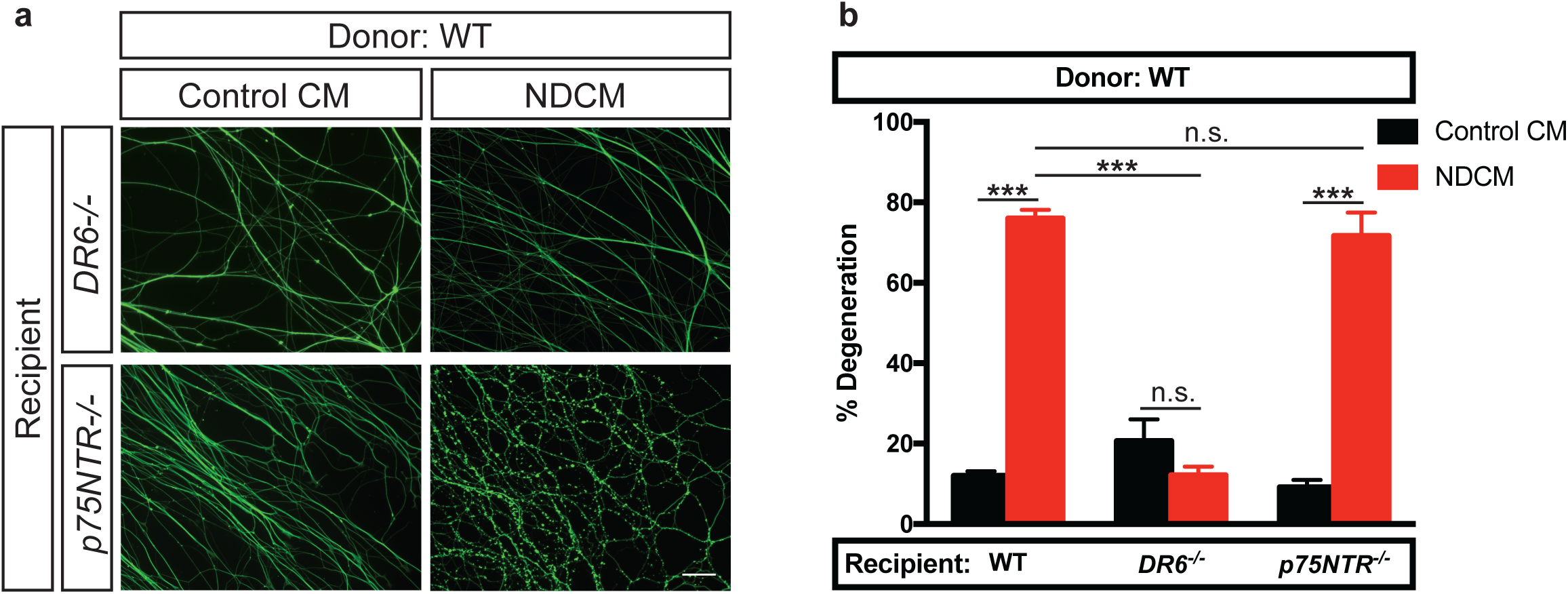
DR6 is required for catastrophic degeneration induced by NDCM. a. Representative images of distal sympathetic axons from *DR6*^*-/-*^ and *p75NTR*^*-/-*^ animals immuno-stained for β3-tubulin after treatment with NDCM and Control CM collected from wild-type neurons for 5 hours, respectively. All cultures were NGF deprived for 12 hours prior to the addition of CM. Scale bar = 50 µm. b. Quantification of (a). Values are represented as mean±SEM. Left two columns represent percentages of degeneration of wild-type sympathetic axons treated with wild-type NDCM and Control CM, respectively. Significant difference is determined by two-way ANOVA with multiple comparisons, ***p<0.0001, n.s.=not significant, n=3 or more for each group. For each repeat, at least 100 axons were scored for degeneration.

## Discussion

By investigating calcium dynamics during the transition from latent to catastrophic phases of degeneration after trophic withdrawal, we identified several novel events: **1.** Intra-axonal calcium rise and spheroid formation between latent and catastrophic phases of degeneration. This spheroid formation is triggered by p75NTR-Rho signaling, caspase activation and actin remodeling. We suggest that the timing of p75NTR-Rho-actin signaling and upregulation of pro-degenerative transcriptional programs may define the duration of the latency phase. **2.** The membranes of growing spheroids rupture leading to expulsion of axoplasmic material to the extracellular environment. This may represent a novel mechanism by which pro-degenerative molecules are released to influence neighboring cells. **3.** The material expelled from ruptured spheroids triggers catastrophic degeneration. We find that DR6 is required to respond to this prodegenerative signal thereby gating entry into the catastrophic phase of degeneration (Fig.8).

**Figure 8:**
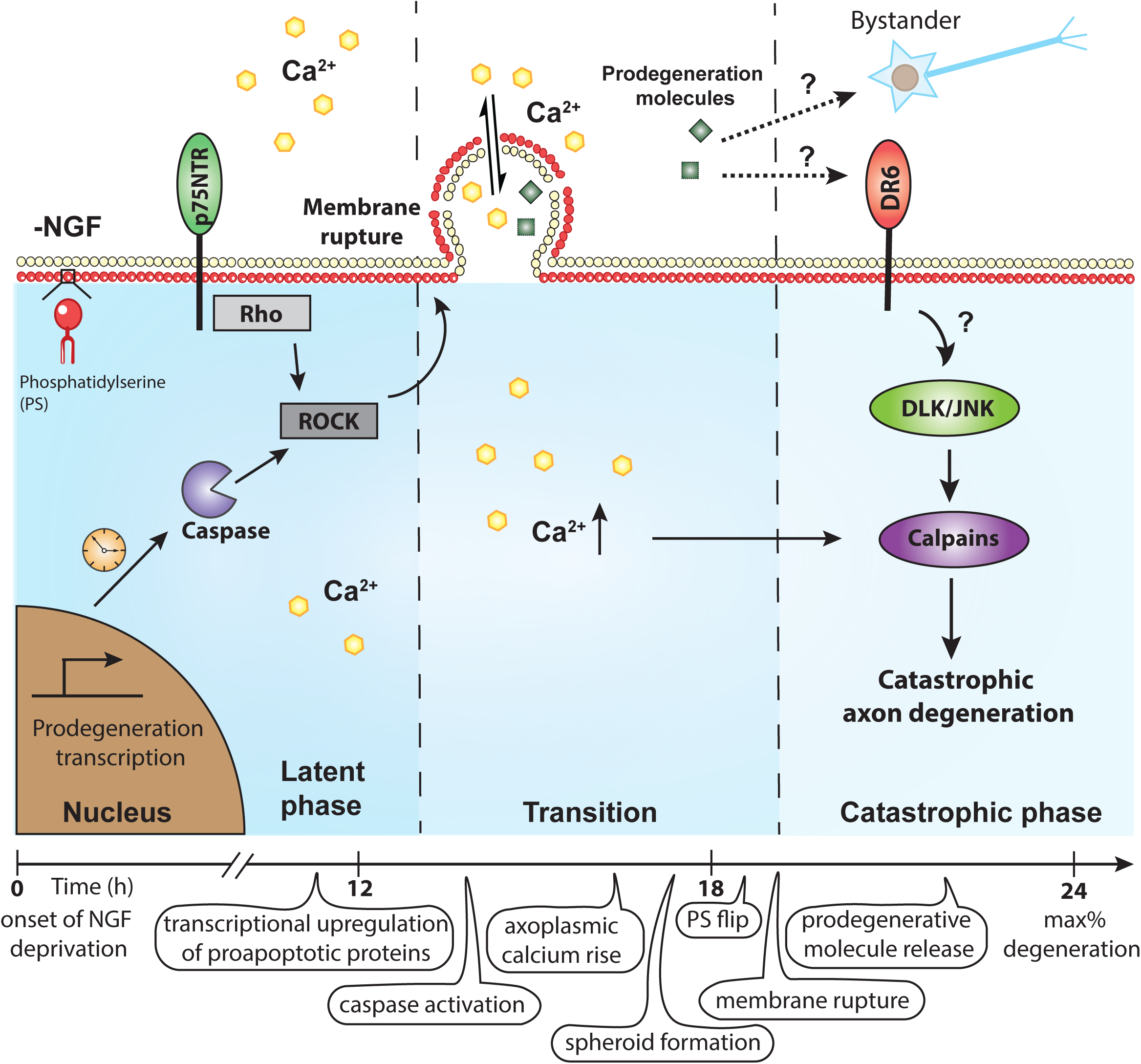
Time course of events associated with NGF-deprivation induced axon degeneration of sympathetic neurons. After NGF deprivation, prodegenerative transcription is upregulated. Axoplasmic calcium is increased and enriched in spheroids prior to catastrophic phase. Spheroid formation is regulated by p75NTR, Rho activity, and caspase activation. The calcium electrochemical gradient across the membrane is disrupted by spheroidal rupture, which may also lead to the release of intra-axonal prodegenerative molecules to extracellular space, acting as extrinsic factors to promote degeneration in a paracrine or autocrine manner. We speculate that p75NTR plays important role in calcium dynamics during the latent phase, while DR6 can be activated by prodegenerative NDCM to mediate downstream catastrophic degeneration pathways (e.g. DLK/JNK, calpastatin, calpain).

Formation of axonal spheroids has been reported in many neurological disorders, including Alzheimer’s disease, ALS, and neurocysticercosis (Beirowski et al., 2010; Griffin and Watson, 1988; Mejia Maza et al., 2018). After injury, spheroids arise along the length of axons. Electron microscopy and immunohistochemistry have revealed that some of the spheroids are empty, while others accumulate mitochondria, lysosomes, autophagosomes, amyloid precursor protein (APP), calpain-mediated spectrin degradation products, and other double membrane and electron dense vesicles (Beirowski et al., 2010). Importantly, calcium overload in these spheroids has been observed on H_2_O_2_ treated axons (Barsukova et al., 2012). While spheroids have been extensively described in the literature, whether they have a function in promoting degeneration has remained unknown. We propose that the function of these spheroids is to trigger catastrophic degeneration by releasing pro-degenerative molecules into the extracellular environment.

Beyond what has been previously described for spheroids (e.g. loaded with mitochondria, ER, and actin), we also report localized annexin V positivity (i.e. PS flipping) (Fig.2c), which suggests they are marked for phagocytic engulfment reminiscent of apoptotic bodies (Segawa et al., 2014; Ravichandran, 2010). Indeed, we observe several instances of spheroids detaching from the axon, which is a unit that presumably would be phagocytosed (Supplemental Movie.2b). In correlation with the growth of spheroids, we also observe that 3kDa and 10kDa fluorescent dextran entering the cytosol consistent with disrupted or ruptured plasma membrane (Fig.3b-c). It is unclear where membrane integrity is disrupted; it may be at the belly of the spheroid and/or at axon/spheroid junctions. Whether or not spheroid rupture also occurs in aforementioned neurodegenerative disorders remains an open question.

We found that disrupting caspase activation, p75NTR, Rho or actin polymerization delays spheroid formation after NGF deprivation (Fig.2a-b, 6b-c). It has previously been shown that the intracellular domain (ICD) of p75NTR modulates RhoA activation by interacting with Rho GDP-dissociation inhibitor (RhoGDI) to inhibit axonal outgrowth and promote axon pruning in response to cues such as myelin-associated glycoprotein (MAG) or oligodendrocyte glycoprotein (OMpg) (Yamashita et al., 2002; Park et al., 2010; Yamashita et al., 1999). In the presence of NGF, the interaction of p75NTR with RhoGDI-RhoA complex is disrupted, leading to RhoA inactivation and neurite outgrowth (Mathew et al., 2009). Consistent with these findings, we show that activation of RhoA in the absence of *p75NTR* promotes entry into catastrophic phase by triggering spheroid formation (Fig.5f-i).

Proneurotrophin activation of p75NTR in the absence of Trk signaling leads to caspase mediated neuronal death (Troy et al., 2002; Gentry et al., 2004). Sublethal executioner caspase activity plays an important role in nervous system development (Unsain and Barker, 2015). Interestingly, inhibition of caspases suppressed spheroid formation, but failed to protect axons from NDCM induced catastrophic degeneration (Fig.4f-g). Thus, in the absence of NGF, spheroid formation is induced by p75NTR, caspase, RhoA and actin remodeling, which are all requisite for transitioning from latent to catastrophic degeneration. It’s unclear whether these factors act in the same pathway or function independently.

Previous work has shown that activation of ROCK1 by caspase-3-mediated cleavage regulates membrane blebbing, independent on Rho activity (Sebbagh et al., 2001; Zhang et al., 2018). Interestingly, recent studies show that trophic deprivation causes a rapid disassembly of the actin-spectrin-based membrane-associated periodic skeleton (MPS), which occurs prior to protein loss and independently of caspase activation (Zhong et al., 2014; Wang et al., 2019). Thus, it’s possible that p75NTR-Rho-actin and caspase-ROCK1 signaling are two independent pathways that modulate spheroid formation at the end of the latent phase of degeneration. Calcium accumulation was accompanied with focal spheroid formation. But the precise relationship between calcium dysregulation, development of spheroids, and p75NTR is not known. Calcium-dependent changes in the cytoskeleton may contribute to spheroid formation as suggested by the appearance of disorganized F-actin when disrupting calcium homeostasis (Barsukova et al., 2012; Oertner and Matus, 2005; Meeker et al., 2016). However, depleting intracellular calcium failed to suppress spheroid formation (Supplementary Fig.2a), indicating that intra-axonal calcium increase after NGF deprivation is not be required to promote spheroid formation.

Our findings are consistent with the notion that p75NTR controls the window of latency prior to catastrophic degeneration. What is the molecular basis for the putative timer controlling the window of latency after trophic withdrawal? Classically, biological timers have been associated with transcriptional programs that form feedback loops (Mitrophanov and Groisman, 2008; Santos and Ferrell, 2008). Although the rates of RNA and protein synthesis decline rapidly in NGF-deprived sympathetic neurons, the expression of a few “death” genes like *c-jun, cyclin D1, Bim, DP6, SM-20* do increase and are critical for axon degeneration (Deshmukh and Johnson, 1997; Maor-Nof et al., 2016; Kristiansen and Ham, 2014; Kristiansen et al., 2011; Freeman et al., 2004). Following NGF withdrawal, JNK activity has been observed to increase over time, resulting in activation of the transcription factor c-Jun in sympathetic neurons (Virdee et al., 1997). Inhibition of transcription by applying actinomycin D to the soma during NGF deprivation blocked spheroid formation and degeneration (Fig.2a-b), indicating the involvement of a degenerative transcriptional program. Consistent with the notion of a transcriptional feedback loop, this pathway has been shown to up-regulate TNFα-converting enzyme (TACE), which is necessary for p75NTR cleavage (Kenchappa et al., 2010). Liberation of the p75NTR ICD retrograde degenerative signal is accompanied by secondary activation of JNK and up-regulation of proapoptotic factors like p53-upregulated modulator of apoptosis (Puma) (Kenchappa et al., 2010; Simon et al., 2016; Pathak et al., 2018). The time it takes to engage this transcriptional program and cleave p75NTR by TACE and γ-secretase may represent a molecular timer governing the duration of the latency window (Pathak and Carter, 2017). This molecular timer may account for the delay of spheroid formation in NGF deprived *p75NTR*^*-/-*^ axons (Fig.5c).

Similar to p75NTR, DR6 is required for axon degeneration after trophic withdrawal (Olsen et al., 2014; Gamage et al., 2017). Does DR6 regulate the same signaling events as p75NTR after trophic factor deprivation? We conclude that this is unlikely given that these TNFR family members govern distinct phases of degeneration. Unlike *p75NTR*^*-/-*^, *DR6*^*-/-*^ axons are capable of expelling prodegenerative axoplasmic materials to the surrounding environment via membrane rupture after NGF deprivation (Fig.5g-h) and *DR6*^*-/-*^ axons are protected from entry into catastrophic phase in response to a 5 hour incubation of NDCM (Fig.7). These findings suggest that DR6 is downstream of the destructive factor(s) found in NDCM, however whether this is a direct or indirect interaction between these factors and DR6 remains to be determined. It’s also unknown how DR6 is activated to promote catastrophic degradation in response to trophic deprivation. DR6 was previously shown to interact with β-amyloid precursor protein (APP) to regulate developmental axonal pruning and synapse restriction, but unlikely to contribute to Alzheimer’s disease related pathology (Nikolaev et al., 2009; Kallop et al., 2014). Given the fact that depletion of APP could partially protect sensory axons from degeneration after NGF withdrawal (Olsen et al., 2014), and that APP isoform expression is altered in the SCG following trophic deprivation (Smith et al., 1993), it’s possible that DR6 promotes developmental catastrophic degeneration via an APP dependent pathway.

Previous findings show that DR6 potently activates NF-κB and JNK to induce apoptosis (Benschop et al., 2009; Hu et al., 2014). Consistent with this, loss of *DR6* suppressed phosphorylation of JNK after injury (Gamage et al., 2017). Thus, we speculate that direct or indirect activation of DR6 by the prodegenerative factor emanating from ruptured spheroids may promote the activation of JNK and calpain, which are known to trigger catastrophic degeneration (Fig.8). It is also possible that activation of DR6 independently activates catastrophic degeneration by engagement of receptor mediated caspase pathways.

While the calcium dependent protease, calpain, is known to be required for catastrophic degeneration, it is unclear why the initial intracellular wave of calcium prior to catastrophic degeneration is insufficient to trigger this event. One clue may come from Tessier Lavigne and colleagues who suggest that depletion of the calpain inhibitor, calpastatin, is required for catastrophic degeneration (Yang et al., 2013). However, the membrane ruptures that we observe at the onset of catastrophic degeneration is likely to cause equilibration of external and internal calcium concentrations. Post-rupture, intraxonal calcium levels are likely to be persistently high, however, depletion of calpastatin is likely prerequisite for calcium dependent catastrophic degeneration. Recent studies show that chelation of extracellular calcium by EGTA in late but not early phases of degeneration rescues axons from trophic deprivation induced degeneration (Johnstone et al., 2019). And the transient receptor potential vanilloid family member 1 (TRPV1) cation channel is required for calcium influx to promote developmental sensory axon degeneration, while plasma membrane nanoruptures allow entry of extracellular calcium to drive axon degeneration in an EAE model of MS (Witte et al., 2019; Johnstone et al., 2019). Thus, extracellular calcium may enter the axoplasma via calcium channels like TRPV1 or spheroidal ruptures. Both routes of sustained intracellular calcium elevation converge on calpain to trigger catastrophic degeneration.

While extensive neuronal apoptosis occurs during development, selective pruning of axons is important for refinement and plasticity of neuronal network. Global and local NGF deprivation in sympathetic neurons cultured in microfluidic devices represent good *in vitro* models to study overlapping and distinct signaling events between these two self-destruction programs. Time-lapse calcium imaging on globally and locally NGF deprived axons reveal similar pre-catastrophic axoplasmic calcium flux (Supplementary Fig.1d-e), indicating a shared pathway to trigger degeneration in late phase. Studies have shown that key mediators like Apaf-1 and caspase 6 are divergent points in axonal apoptosis and pruning pathways (Cusack et al., 2013). Whether modulating these divergent factors show different calcium dynamics and spheroid pattern during axon degeneration requires further investigation.

Our data reveal novel signaling emanating from two different death receptors to govern latent and catastrophic phases of degeneration. After NGF deprivation, prodegenerative transcription, caspase activation and p75NTR-Rho dependent actin remodeling promote formation of axonal spheroids. Axoplasmic materials containing prodegenerative molecules are then released to the extracellular space via membrane rupture, which may act as a positive feedback loop to hasten the entry of axons into catastrophic phase degeneration(Fig.8). Entry into the catastrophic phase of degeneration appears to be gated by DR6 (Fig.8). Whether or not this represents a mechanism whereby axons might coordinate their degeneration requires further study. Since axonal spheroids have been described in many neurological disorders and injury models, the destructive role of spheroids may extrapolate to other degenerative etiologies. For example, intracellular toxic aggregated proteins like tau proteins could be released into extracellular space via spheroidal ruptures, which contributes to the pathogenesis of neurodegenerative disease. Thus, the notion of destructive spheroids and subsequent inter-axonal communication during degeneration facilitates our understanding of neural refinement during development and presents intriguing possibilities with respect to therapeutic intervention to alleviate bystander degeneration in disease.

## Materials and Methods

### Mice

All experiments were carried out in compliance with the Association for Assessment of Laboratory Animal Care policies and approved by the University of Virginia Animal Care and Use Committee. All mice are on a C57BL/6J.129S mixed background. *p75NTR*^*-/-*^ animals were purchased from the Jackson labs. *DR6*^*-/-*^ animals were a generous gift from the Genentech. Males and females are mixed in all of our experiments.

### Primary Sympathetic neuronal cultures

Sympathetic neuron cultures were established as described previously (Deppmann et al., 2008). Briefly, neurons were obtained by dissociation of P0-P2 mouse superior cervical ganglia. These neurons (from each litter of pups) were plated in compartmentalized microfluidic devices in DMEM supplemented with 10% FBS, penicillin/streptomycin (1U/mL), and 45 ng/mL of NGF purified from mouse salivary glands. Glia are removed from cultures using 5 μM cytosine arabinofuranoside (Ara-C) (Sigma) for 48-72 hours. For NGF deprivation, cultures were washed three times with medium lacking NGF and then maintained in NGF-deficient media containing a neutralizing antibody (1μg/mL anti-NGF antibody, Millipore) through designated time points at 37°C.

### Fabrication and use of microfluidic devices

Microfluidic devices were generated as described previously (Park et al., 2006). These chambers were affixed to coverglass coated with poly-D-lysine (50μg/mL) and laminin (1μg/mL).

### Immunocytochemistry

Immunocytochemistry was carried out as previously described (Singh et al., 2008). Briefly, at indicated times, axons were fixed in 4% paraformaldehyde (w/v)/ phosphate buffered saline (PBS) at room temperature for 20 minutes, washed 3×5min with 1x PBS, and blocked/permeabilized (5% goat serum, 0.05% Triton-x-100 in PBS) for 1 hour at room temperature. Axons were then incubated overnight at 4°C with primary antibody diluted in blocking buffer. Cells were then washed 3×5 min with 1x PBS and incubated with fluorescent secondary antibody for 1 hour at room temperature. Cells were again washed with 1x PBS three times and imaged using a fluorescent inverted microscope. The antibodies used in this study are mouse anti-Tuj1 (1:1000, Covance), and goat anti-mouse Alexa 488 (1:800, Life Technologies). For active Rho staining, axons were incubated with 50μg/mL of GST-Rhotekin-RBD fusion protein in blocking solution for 1 hour at room temperature, washed 3×5min with 1x PBS. Cells were then incubated with anti-GST (1:800, Sigma G1160) diluted in blocking buffer for 1 hour, washed 3×5min with 1x PBS and incubated with fluorescent secondary antibody for 1 hour at room temperature. All *in vitro* experiments were performed in triplicate with at least two microfluidic devices used for each condition.

### Live imaging

Sympathetic neuron cultures were washed 3 times with DMEM/F-12, Phenol Red free, and incubated for 30 minutes at 37°C and 10% CO_2_ with live imaging dyes diluted in DMEM/F-12, Phenol Red free. Cells were then imaged under Leica SP5 X confocal microscope in W.M. Keck Center in University of Virginia. Axons in grooves of microfluidic chamber were imaged after NGF deprivation. For assessing flipping of phosphatidylserine on axonal spheroids, Annexin V red reagent (IncuCyte, 4641) diluted in DMEM/F-12, Phenol Red free (1:200) after NGF deprivation. For membrane rupture, dextran dyes diluted in DMEM/F-12, Phenol Red free were added to the microfluidic chamber after 17 hours NGF deprivation. The dyes used in this study are Fluo-4 AM (1μM, F14201) and Dextran Texas Red, neutral, 3 kDa (50μM, D3329), 10 kDa (50μM, D1828) and 70 kDa (50μM, D1830). All dyes were purchased from Thermo Fisher Scientific.

### Image Processing and analysis

Axon degeneration in culture was quantified from β3-tubulin stained fluorescence images by counting the number of individual axons at the leading edge that had at least three beads/blebs as described (Zhai et al., 2003). A blinded investigator counted ten representative pictures of the axons, in two microfluidic chambers per condition/time point. On each image 10 50μm boxes were randomly assigned to single axons. The investigator took care not to box bundles of axons, which may confound analysis. Then the number of boxes, which had 3 or more beads/blebs were counted and categorized as degenerating axons. Equal to or more than 80% degeneration was considered maximum degeneration and equal to or less than 10% degeneration of axons was considered as minimum degeneration. The percentage of the total number of degenerating axons was calculated using Microsoft Excel. At least 300 total axons were counted for each condition. The standard error of the mean was considered as error. In live imaging, Ca^2+^ intensity (ΔF/F0), the size (S/S0) and number of axonal spheroids were quantified in selected ROI (single axon or axonal spheroid) by Fiji software (Schindelin et al., 2012). Each experiment was repeated at least 3 times with separate litters of mouse pups of the same genotype.

### Conditioned media experiments

Established compartmentalized neuron cultures were deprived of NGF by washing with NGF free, serum free media three times and incubating at 37°C with a neutralizing anti-NGF antibody for 12 hours. NGF deprivation conditioned media (NDCM) was collected from the axonal chamber of the device after incubating at 37°C with a neutralizing anti-NGF antibody for 24 hours. A volume of ∼100μL of NDCM was pooled per device from more than 20 devices per experiment. The media in the wells of the device on the axonal side was also collected. Importantly, fresh serum free/NGF free media is added to distal axons before incubation so as not to confound analysis with factors secreted during normal growth. NDCM or control conditioned media was applied on axons for 5 hours, which were pre-grown in the absence of NGF for 12 hours.

### Calcium measurement

Conditioned media was diluted to milliQ water (1:20) and mixed thoroughly. 100μL of reaction mixture was made with 1μL HEPES, 2μL 1mM Fluo-4 (20mM working concentration, Life Technologies), 10μL diluted conditioned media, and 87μL water. Black 96 well plate was used in Spectrophotometric assay. Eight CaCl_2_ standards (2.54μM, 4.87μM, 9.75μM, 19.5μM, 39μM, 78μM, 156μM, and 313μM) were used to calculate standard curve for analyzing Ca^2+^ concentration in conditioned media.

### Exogenous Calcium assay

Sympathetic neuron cultures were washed three times with DMEM supplemented with 10% FBS, penicillin/streptomycin (1U/mL), and then incubated in NGF deficient media containing a neutralizing antibody (1μg/mL anti NGF antibody, Millipore) for 12 hours. For exogenous calcium concentration below 1.8mM (DMEM), cells were washed 3 times with DMEM Ca^2+^ free media. CaCl_2_ was added to DMEM Ca^2+^ free media to indicated final concentrations, which was them applied to cultures 37°C and 10% CO_2_ for 5 hours.

### Statistical Analysis

Statistical analysis was performed in graphpad prism software and Excel as indicated in figure legends. All data shown represents the mean ± SEM. Sample number (n) was defined as the number of individuals that were quantified in each experiment. The level of significance (α) was always set to 0.05. For ANOVA, if statistical significance was achieved, we performed post hoc analysis to account for multiple comparisons.

## Acknowledgements

We thank Pamela Neff, Elena Tenore, Nadine Ly, and Courtney Cushman for technical assistance. Austin Keeler, Sushanth Kumar, Shayla Clark, Amrita Pathak, Vitaly Zimyanin and Brandon Podyma for helpful discussions and comments on the manuscript. We thank Xiaorong Liu, Barry Condron, and Ali Guler for helpful comments on the manuscript. We thank Ammasi Periasamy and acknowledge the Keck Center for Cellular Imaging for the usage of the Leica SP5X microscopy system (RR025616). We wish to thank Genentech for kindly providing the *DR6*^*-/-*^ mouse line. This work was supported by NIH-NINDS grant R01NS091617 and the Owens family foundation awarded to C.D.D.

## Supplementary Figures

**Supplementary Figure 1: Calcium dynamics in response to global and local NGF deprivation**

a. Schematic representation of local NGF deprivation paradigm in microfluidic devices. Cell bodies (CB) and distal axons (DA) are separated. For local NGF deprivation, sympathetic distal axons were maintained in 80µL per well of NGF-deficient media containing 1µg/mL anti-NGF antibody. Cell bodies were maintained in 150µL per well of media containing 45ng/mL NGF.

b. Representative images of β3-tubulin immuno-stained distal sympathetic axons before treatment (0hr), 12, 24, 36, 48, and 72 hours after local NGF deprivation. Scale bar = 50µm.

c. Degeneration time course after local NGF deprivation. Nonlinear regression curves drawn according to Hill equation. n=3 for each time point. Data points represent as mean±SEM.

d. Normalized calcium fluorescence change of global NGF deprived axons over time (0-27 hours). Green line represents the mean of intra-axonal calcium fluorescent change. Total number of n=6 axons from 3 independent litters were quantified. Data shown as mean±SEM.

e. Normalized calcium fluorescence change of local NGF deprived axons over time (36-54 hours). Green line represents the mean of intra-axonal calcium fluorescent change. Total number of n=11 axons from 3 independent litters were quantified. Data shown as mean±SEM.

f. Fluo4-AM calcium imaging of sympathetic axons at indicated times in the presence of NGF. Scale bar = 10µm.

g. Quantification of normalized calcium fluorescence of axonal spheroids 30 to 90 minutes after 17 hours of global NGF deprivation. Individual axonal spheroids were quantified: n=18 spheroids from n=3 independent replicates. Data shown as mean±SEM.

**Supplementary Figure 2: Axonal spheroid formation and membrane disruption after NGF deprivation**

a. Fluo4-AM calcium imaging of sympathetic axons after 18 hours of NGF deprivation with DMSO or intracellular calcium chelator BAPTA-AM (10µM, 1hr incubation before NGF deprivation). Scale bar = 10µm. Number of spheroids per 100µm of axon after 18 hours of NGF deprivation were quantified on the right. Individual axons were counted: n=25 (DMSO) and n=26 (BAPTA-AM) axons from 3 independent replicates. Significance is determined by unpaired t-test, n.s.=not significant. Data shown as mean±SEM.

b. Phase contrast imaging of axonal spheroids after 18 hours of NGF deprivation. Scale bar = 5µm.

c. Representative images of axonal spheroids after 18 hours of NGF deprivation. Annexin V (red) indicates exposure of phosphatidylserine on the extracellular surface of axonal spheroids.

d-e. Examples of individual spheroidal tracing on NGF deprived axons labeled with Fluo4-AM and bathed in 3 kDa (c) and 10 kDa (b) dextrans, respectively. Right y-axis is the mean grey level intensity of dextran fluorescence (red) after background subtraction, while left y-axis is the mean grey level intensity of Fluo4-AM (green) after background subtraction. Spheroids showing the red/green transition are captured.

**Supplementary Figure 3: Elevated calcium concentration in NDCM**

a. Schematic representation of the spectrophotometric assay using Fluo4 (cell impermeant) to measure calcium concentration in conditioned media. The standard curve showing the relative fluorescence units (RFU) as a function of increasing calcium concentration.

b. Measurement of extracellular calcium concentration of untreated and NGF deprived conditioned media. In “Control CM” group, media was collected from untreated axons in the presence of NGF. In “NDCM” groups, medium were collected from axons after 12, 18, and 24 hours of NGF deprivation, respectively. Values were analyzed from n=7 (Control CM), n=4 (12hrs NDCM), n=6 (18hrs NDCM), and n=8 (24hrs NDCM) independent replicates. Significance is determined by one-way ANOVA with multiple comparisons, *p<0.05. Data shown as mean±SEM.

c. Measurement of extracellular calcium concentrations in Control CM and 18hrs NDCM in the absence and presence of NCX blocker 10µM Bepridril or PMCA blocker 0.5mM Caloxin 2A1. Values were analyzed from n=4 (Control CM), n=4 (18hrs NDCM, --), n=3 (18hrs NDCM, Bepridril), and n=3 (18hrs NDCM, Caloxin 2A1) independent replicates. Significance is determined by one-way ANOVA with multiple comparisons, *p<0.05. Data shown as mean±SEM.

d. Measurement of extracellular calcium concentration of untreated and NGF deprived conditioned media in the presence and absence of intracellular calcium chelator BAPTA-AM (10µM, 1hr incubation before NGF deprivation), SERCA inhibitor Thapsigargin (100nM, overnight treatment before NGF deprivation), and mitochondrial mPTP blocker Cyclosporin A (20µM, 1 hour treatment before NGF deprivation). Calcium concentrations were measured from at least 3 independent experiments. Significance is determined by two-way ANOVA with multiple comparisons, n.s.=not significant, *p<0.05. Data shown as mean±SEM.

e. Measurement of extracellular calcium concentration of +NGF (Control CM) and 18 hours NGF deprived conditioned media (18hrs NDCM) collected from wild-type, *DR6*^*-/-*^ and *p75NTR*^*-/-*^ sympathetic neurons. Values were analyzed from n=7 (Control CM, WT), n=6 (18hrs NDCM, WT), n=6 (Control CM, *DR6*^*-/-*^), n=5 (18hrs NDCM, *DR6*^*-/-*^), n=4 (Control CM, *p75NTR*^*-/-*^), n=5 (18hrs NDCM, *p75NTR*^*-/-*^) independent replicates. Significant difference is determined by two-way ANOVA with multiple comparisons, n.s.=not significant, **p<0.001, ***p<0.0001. Data shown as mean±SEM.

**Supplementary Figure 4: Spheroid formation induced by trophic deprivation requires p75NTR, Rho activity and actin remodeling**

a. Representative axons/spheroids visualized for β3-tubulin (Tuj1), Phalloidin and DIC after 18 hours of NGF deprivation. Scale bar = 5 µm.

b. Quantification of axonal spheroid number per 100µm of wild-type sympathetic axon at indicated times after NGF deprivation in the absence and presence of Cytochalasin D or CT04. Total number of n=9 (Control), n=22 (Cytochalasin D), and n=8 (CT04) axons from 3 independent litters were quantified. Values are represented as mean±SEM. Significance is determined by two-way ANOVA with multiple comparisons, ***p<0.0001.

c. Quantification of axonal spheroid number per 100µm of *p75NTR*^*-/-*^ sympathetic axon at indicated times after NGF deprivation in the absence and presence of CN03. Individual axons were counted: n=16 (Control) and n=12 (CN03) axons from 3 independent replicates. Values are represented as mean±SEM. Significance is determined by two-way ANOVA with multiple comparisons, ***p<0.0001.

d. Measurement of extracellular calcium concentration of untreated and NGF deprived conditioned media in the presence and absence of CT04 or CN03 collected from wild-type (black) and *p75NTR*^*-/-*^ (blue) sympathetic axons. Calcium concentrations were measured from at least 3 independent experiments. Significance is determined by one-way ANOVA with multiple comparisons, n.s.=not significant, *p<0.05, **p<0.001, ***p<0.0001. Data shown as mean±SEM.

e. Measurement of extracellular calcium concentration of untreated (Control CM), 18 hours and 24 hours NGF deprived conditioned media from *p75NTR*^*-/-*^ sympathetic axons. Values were analyzed from n=4 (Control CM), n=5 (18hrs NDCM), and n=3 (24hrs NDCM) independent replicates. Significance is determined by one-way ANOVA with multiple comparisons, n.s.=not significant, *p<0.05. Data shown as mean±SEM.

f. Quantification of degeneration of wild-type and *p75NTR*^*-/-*^ sympathetic with or without 24 hours of NGF deprivation. Significant difference is determined by two-way ANOVA with multiple comparisons, ***p<0.0001, n=3 or more for each group. For each repeat, at least 100 axons were scored for degeneration. Data shown as mean±SEM.

**Supplementary Figure 5: Spheroid formation induced by trophic deprivation requires ligand binding to p75NTR**

a. Fluo4-AM calcium imaging of wild-type and *p75NTR*^*-/-*^ sympathetic axons after 18 hours of NGF deprivation in the absence and presence of 20µg/mL anti-BDNF and 2ng/mL p75NTR ligand/functional blocker LM11A-31. Scale bar = 10µm.

b. Quantification of spheroid number per 100µm of axons in (e). Individual axons were counted: n=15 (WT, Control), n=16 (WT, anti-BDNF), n=37 (WT, LM11A-31), n=16 (*p75NTR*^*-/-*^, Control), n=19 (*p75NTR*^*-/-*^, anti-BDNF), and n=18 (*p75NTR*^*-/-*^, LM11A-31) axons from 3 independent replicates. Significance is determined by two-way ANOVA with multiple comparisons, n.s.=not significant, ***p<0.0001. Data shown as mean±SEM.

**Supplementary movie 1: Axoplasmic calcium dynamics and PS flipping of spheroidal membrane with or without NGF deprivation**

a. Fluo4-AM (green) and Annexin V (red) live imaging of wild-type sympathetic axons with NGF. Scale bar = 10µm.

b. Fluo4-AM (green) and Annexin V (red) live imaging of wild-type sympathetic axons after 18 hours of NGF deprivation. White arrows indicate calcium enriched spheroids with PS flipping. Red arrow indicates ruptured spheroids. Yellow arrowheads indicate calcium enriched spheroids without membrane disruption. Scale bar = 10µm.

**Supplementary movie 2: Spheroidal membrane rupture after NGF deprivation**

a. Live imaging of dextran 3 kDa (red) entry to axonal spheroids (black) after 18 hours of NGF deprivation. Yellow empty arrows indicate that dextran 3 kDa enter to axonal spheroids. Scale bar = 10µm.

b. Live imaging of dextran 10 kDa (red) entry to axonal spheroids (black) after 19.5 hours of NGF deprivation. Yellow empty arrows indicate that dextran 10 kDa enter to axonal spheroids. White filled arrows indicate spheroids detaching from the axon. Note that some spheroids display punctate labeling consistent with macropinocytosis prior to filling, which is consistent with rupture. Scale bar = 10µm.

c. Live imaging of dextran 3 kDa (red) exclusion of sympathetic axons in the presence of NGF. Scale bar = 10µm.

