## Supplementary figures and images for "p75NTR and DR6 regulate distinct phases of axon degeneration demarcated by spheroid rupture"

### supplemental figures

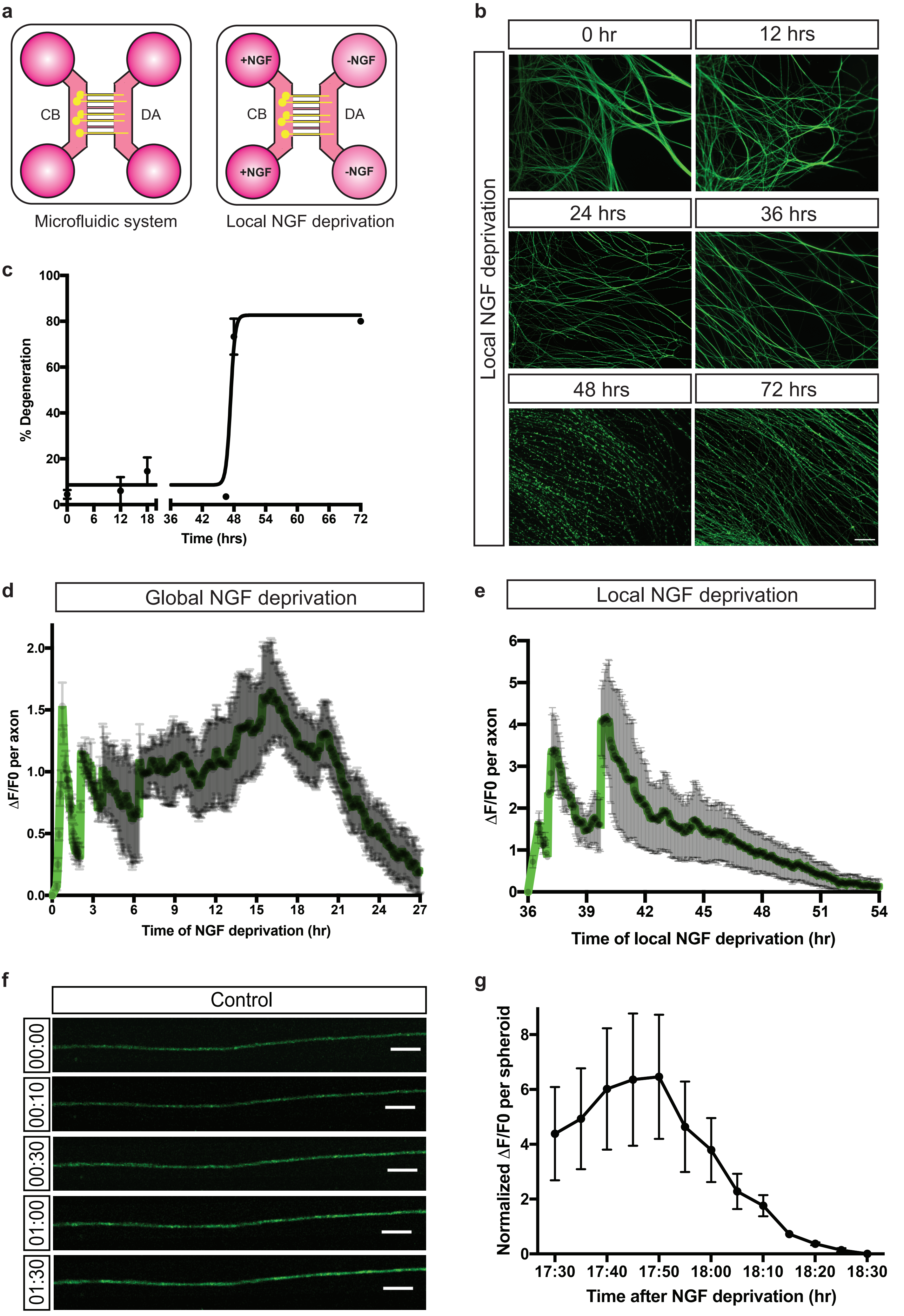


Supplementary Figure 1

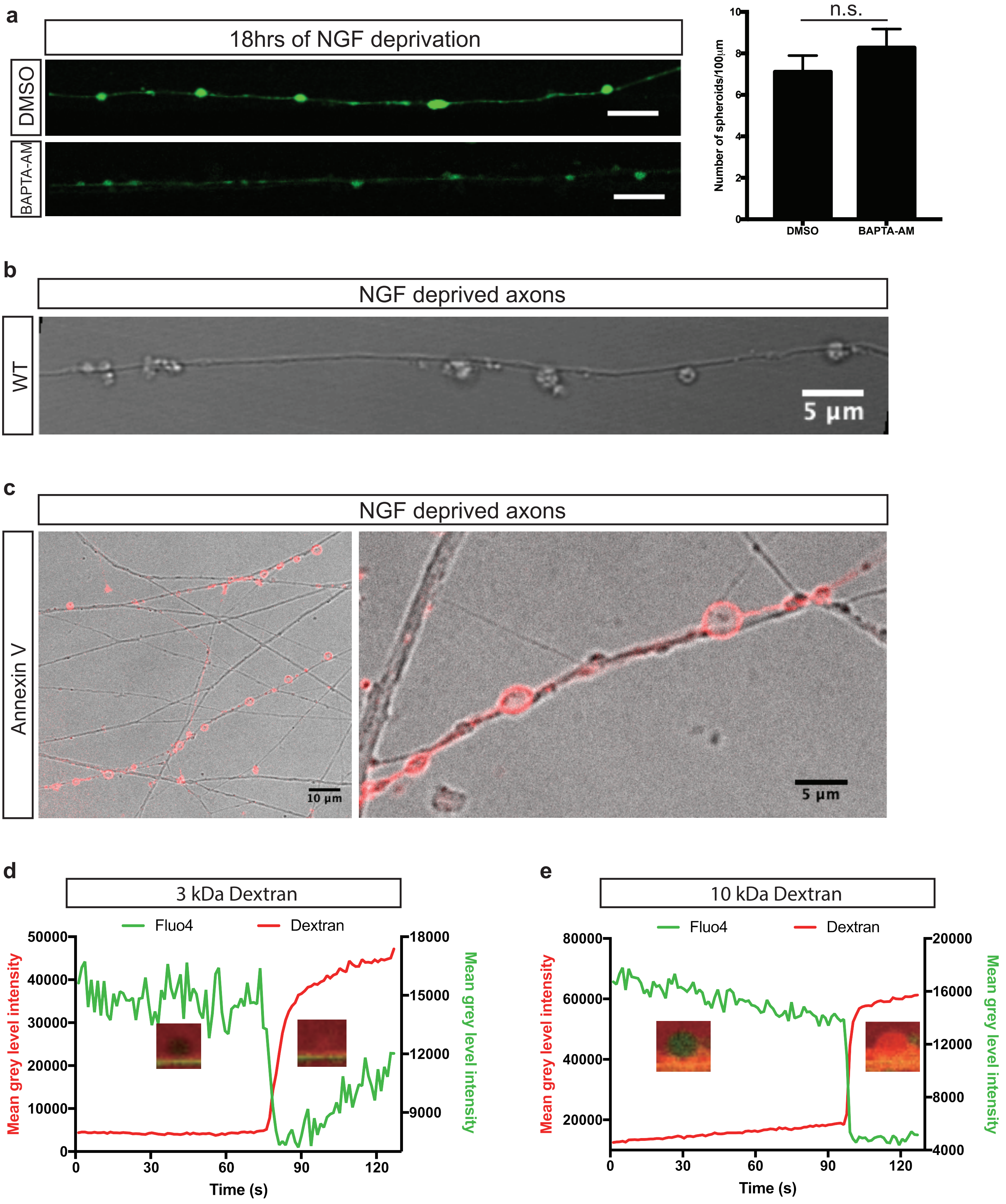

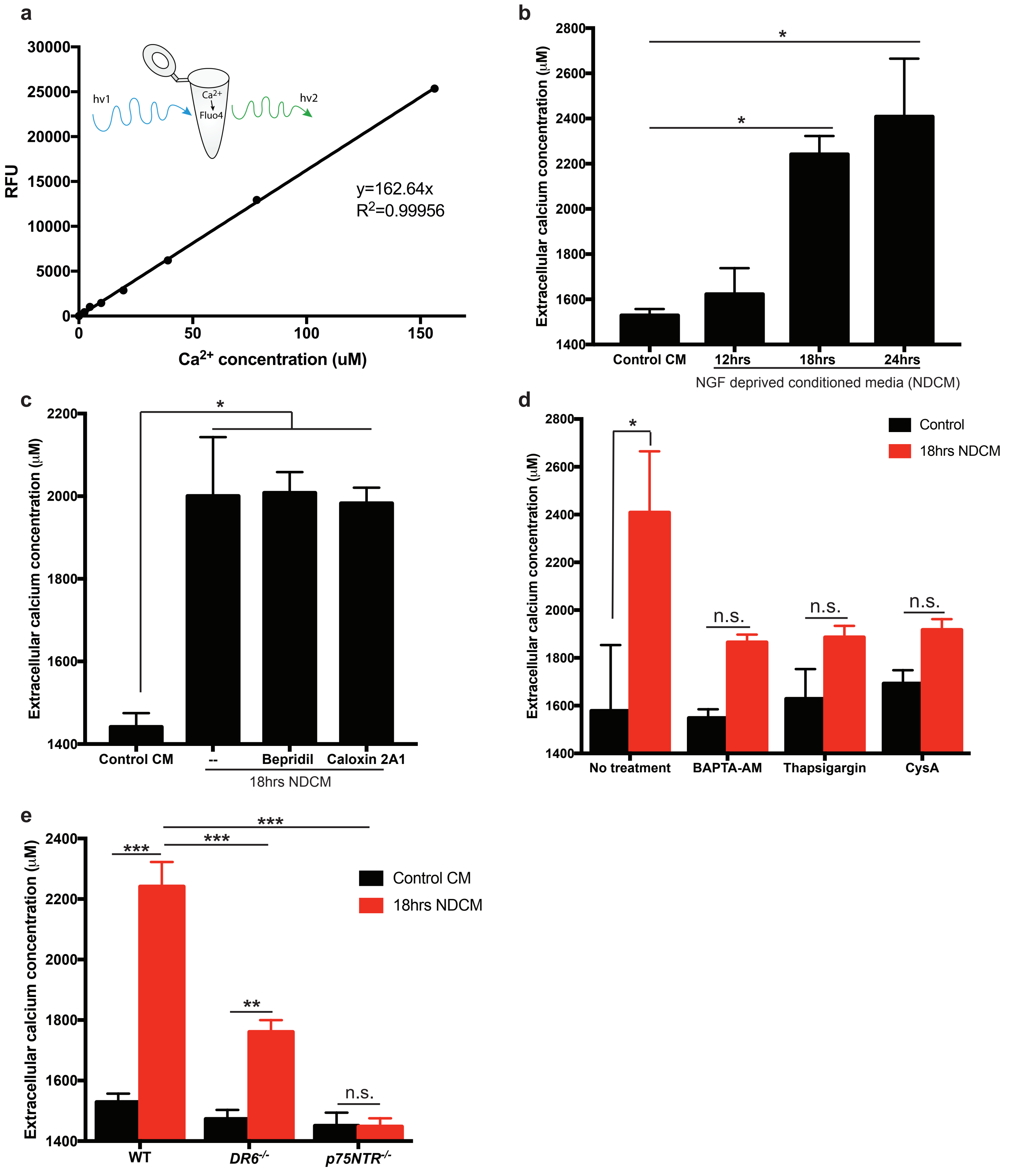


Supplementary Figure 3

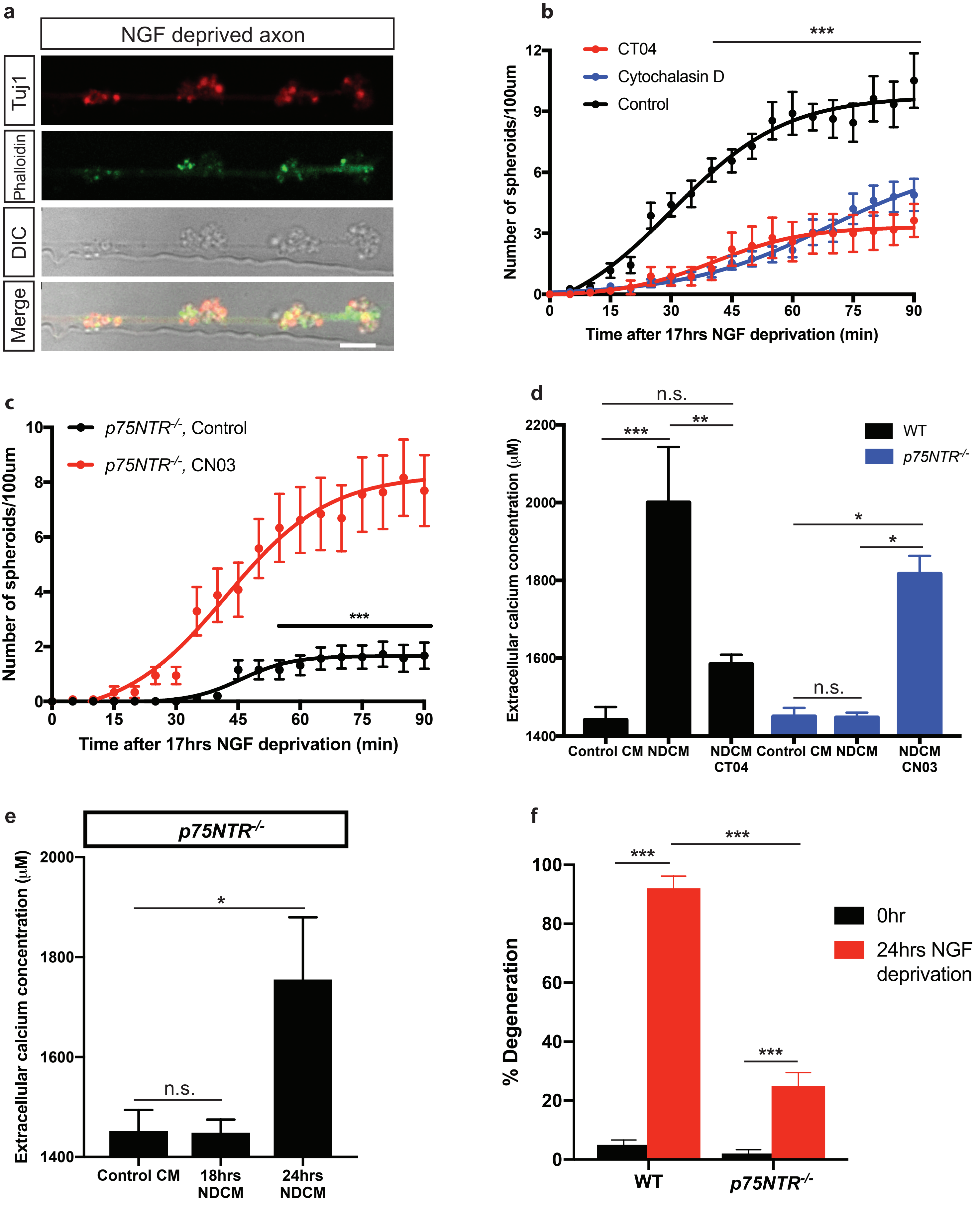


Supplementary Figure 4

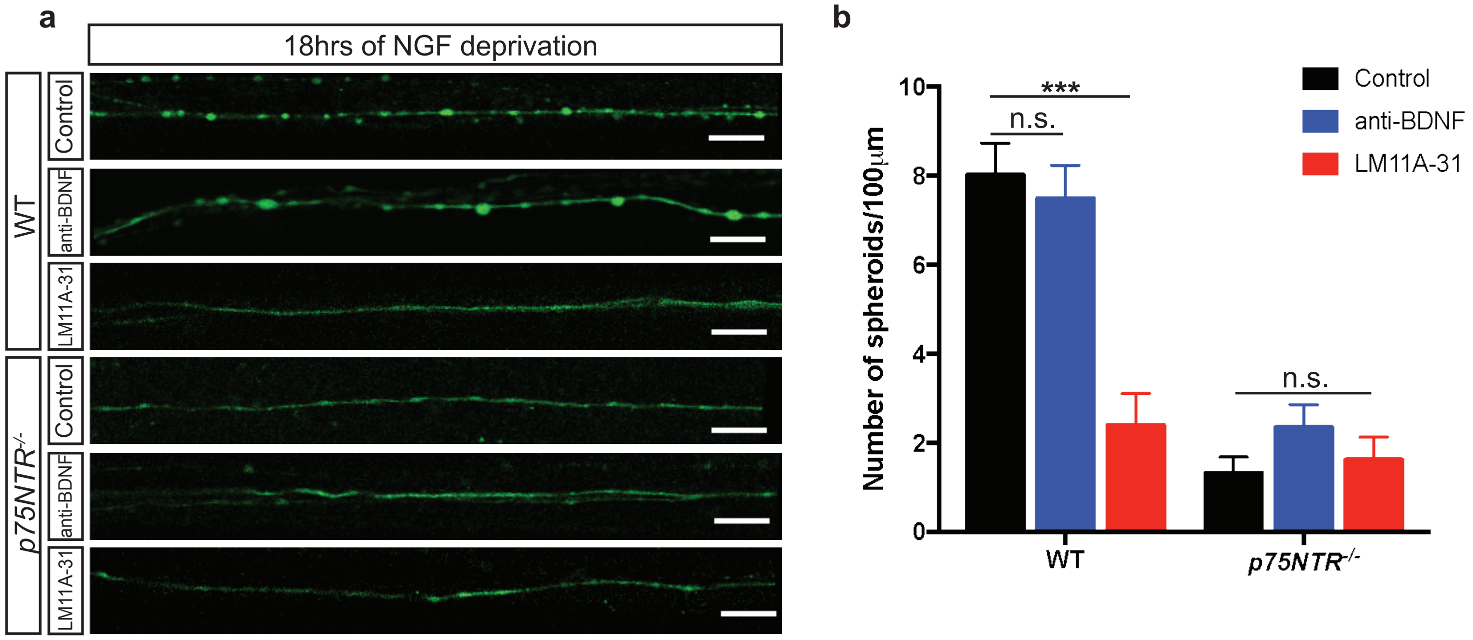
